# The ApiAP2 factor PfAP2-HC is an integral component of heterochromatin in the malaria parasite *Plasmodium falciparum*

**DOI:** 10.1101/2020.11.06.370338

**Authors:** Eilidh Carrington, Roel H. M. Cooijmans, Dominique Keller, Christa G. Toenhake, Richárd Bártfai, Till S. Voss

## Abstract

Malaria parasites undergo a highly complex life cycle in the human host and the mosquito vector. The ApiAP2 family of sequence-specific DNA-binding proteins plays a dominant role in parasite development and life cycle progression. Of the ApiAP2 factors studied to date, most act as transcription factors regulating stage-specific gene expression. Here, we characterised a new ApiAP2 factor in *Plasmodium falciparum* (PF3D7_1456000) that we termed PfAP2-HC. Via detailed investigation of several single or double genetically engineered parasite lines, we demonstrate that PfAP2-HC specifically binds to heterochromatin throughout the genome. Intriguingly, PfAP2-HC does not bind DNA *in vivo* and recruitment of PfAP2-HC to heterochromatin is independent of its DNA-binding domain but strictly dependent on heterochromatin protein 1. Furthermore, our results suggest that PfAP2-HC functions neither in the regulation of gene expression nor in heterochromatin formation or maintenance. In summary, our findings reveal that PfAP2-HC constitutes a core component of heterochromatin in malaria parasites. They furthermore identify unexpected properties of ApiAP2 factors and suggest substantial functional divergence among the members of this important family of regulatory proteins.

## Introduction

The apicomplexan parasite *Plasmodium falciparum* is the main cause of severe malaria worldwide, with the majority of the estimated 405,000 malarial deaths in 2018 attributed to this pathogen ^1^. The symptoms of the disease occur due to repeated asexual intraerythrocytic developmental cycles (IDC), where merozoite stage parasites invade human red blood cells (RBCs) and develop through the ring stage [0-24 hours post-invasion (hpi)] and trophozoite stage (24-30 hpi), before undergoing schizogony to produce mature segmented schizonts containing up to 32 merozoites (30-48 hpi). Rupture of the infected RBCs (iRBCs) releases the merozoites, which in turn undergo another IDC after invading new RBCs. A small proportion of schizonts per cycle commit to the sexual development pathway and produce ring stage daughter cells that mature over a period of ten days and through four intermediate stages (I-IV) into mature stage V gametocytes ^2^. Circulating stage V gametocytes are the only forms of the parasite able to infect the mosquito vector and are therefore essential for malaria transmission.

A key trait of *P. falciparum* is the ability to adapt to and evade the constantly changing environment in its human host through clonally variant gene expression, a process vital to a broad range of biological processes, including antigenic variation, RBC invasion, solute transport and sexual conversion ^3–5^. Clonally variant gene expression in *P. falciparum* is regulated epigenetically, with heritable gene silencing mediated by heterochromatin ^6^. Heterochromatin is found at subtelomeric regions on all 14 chromosomes and in some chromosome internal islands and is characterized by the binding of heterochromatin protein 1 (PfHP1) to the histone modification histone 3 lysine 9 trimethylation (H3K9me3) ^7–11^. These PfHP1/H3K9me3-demarcated heterochromatic domains cover over 400 genes in total (approximately 8% of all protein-coding genes in the genome) ^7,10^. As a core component of heterochromatin, PfHP1 plays an essential role in heterochromatic gene silencing and has a multi-faceted role in parasite biology as previously demonstrated with a conditional loss-of-function mutant ^12^. Conditional depletion of PfHP1 resulted in the de-repression of multi-copy gene families important in antigenic variation, including the well-characterized *var* gene family ^12,13^. In addition, around half of progeny parasites depleted of PfHP1 underwent gametocytogenesis due to de-repression of the internal heterochromatic *pfap2-g* locus encoding the master transcriptional regulator of gametocytogenesis, PfAP2-G, ^12,14,15^. The remaining progeny arrested at the trophozoite stage, indicating an essential role of PfHP1 in proliferation ^12^. With such a diverse range of processes reliant on PfHP1 and heterochromatic gene silencing, the mechanisms of this system warrant further study. However, the molecular machinery involved in heterochromatin establishment, spreading and maintenance in *P. falciparum* remain elusive, along with the transcription factors involved in regulating the expression of heterochromatic genes.

The main transcription factor family in Apicomplexan parasites is the ApiAP2 group of DNA-binding proteins, comprising 27 members in *P. falciparum* ^16–18^. ApiAP2 proteins are characterized by the presence of one to three AP2 domains, homologous to the DNA-binding domains of plant APETALA2/ethylene response element binding protein (AP2/EREBP) transcription factors ^16,19^. To date, five members have been functionally analysed in *P. falciparum*, three of which are acting as transcription factors. PfAP2-G, as mentioned above, is the master regulator of sexual commitment ^14^ and has recently been confirmed as an activator of gametocyte genes, with an additional role in regulating RBC invasion genes suggested ^20^. PfAP2-I is likely essential for parasite survival and regulates a subset of gene families involved in RBC invasion ^21^. Interestingly, PfAP2-I and PfAP2-G also bind upstream of several genes encoding ApiAP2 factors, which could suggest a complex regulatory interplay between ApiAP2 family members ^20,21^. Indeed, Josling and colleagues provided evidence of cooperative binding of PfAP2-G and PfAP2-I to some invasion gene promoters ^20^. PfAP2-EXP is involved in regulating multi-gene families, including *rif*, *stevor* and *pfmc-2tm*, and is seemingly essential for asexual growth ^22^. In contrast, PfSIP2 predominantly binds to SPE2 motifs found in telomere-associated repeat elements (TAREs) and upstream of subtelomeric upsB *var* genes, both of which are heterochromatic, suggesting a possible role in heterochromatin and/or chromosome end biology ^23^. Finally, PfAP2-Tel binds to telomere repeats on all 14 chromosomes and is likely involved in telomere maintenance mechanisms ^24^. Beyond these studies in *P. falciparum*, much has been achieved in characterising ApiAP2 proteins of *Plasmodium* species infecting rodents. In *P. berghei*, several ApiAP2 factors with essential roles in gametocytogenesis ^15,25,26^ and in the mosquito and liver stages ^27–30^ have been studied. Additionally, systematic knockout screens in *P. berghei* ^31^ and *P. yoelii* ^32^ provided an extensive characterisation of the ApiAP2 family and highlight essentiality at different life cycle stages. Interestingly, although some orthologues have the same function in *P. falciparum* and *P.berghei*, for example AP2-G ^14,15^, others display differences such as the PfAP2-EXP orthologue PbAP2-SP, which is expressed exclusively in the sporozoite stages of *P. berghei* ^29^. Interestingly, we recently identified the ApiAP2 protein PF3D7_1456000 as a putative interaction partner of PfHP1 using co-immunoprecipitation (co-IP) experiments coupled with protein mass spectrometry ^33^. Here, we present a multifaceted approach to dissect the potential functions of this ApiAP2 factor in heterochromatin-associated processes during blood stage development of *P. falciparum* parasites.

## Results

### PfAP2-HC specifically associates with heterochromatin

We recently identified a list of potential PfHP1 interaction partners, which includes a member of the ApiAP2 family of putative transcription factors, PF3D7_1456000 ^33^, hereafter referred to as PfAP2-HC. To validate the interaction between PfAP2-HC and PfHP1, we employed a two-plasmid CRISPR/Cas9 based gene editing approach to N-terminally tag PfAP2-HC with GFP (GFP-PfAP2-HC) (Fig. 1a, Supplementary Fig. 1). We tagged the N-terminus because the single AP2 domain of PfAP2-HC is located right at the C-terminus of the protein where tagging may interfere with its function. We obtained a clonal line of the resulting 3D7/GFP-PfAP2-HC population by limiting dilution cloning and confirmed correct editing of the locus by PCR on genomic DNA (gDNA) (Supplementary Fig. 1). Live cell fluorescence imaging of GFP-PfAP2-HC revealed a perinuclear localization, which was undetectable in ring stages and first appeared in trophozoites, peaking mid-schizogony and decreasing in late schizonts (Supplementary Fig. 1). This temporal expression pattern is consistent with the transcriptional profile of *pfap2-hc* during the IDC ^34^. The localisation pattern of GFP-PfAP2-HC matches that of PfHP1 in immunofluorescence assays (IFAs), where the two proteins appear to overlap (Fig. 1b).

**Figure 1.**
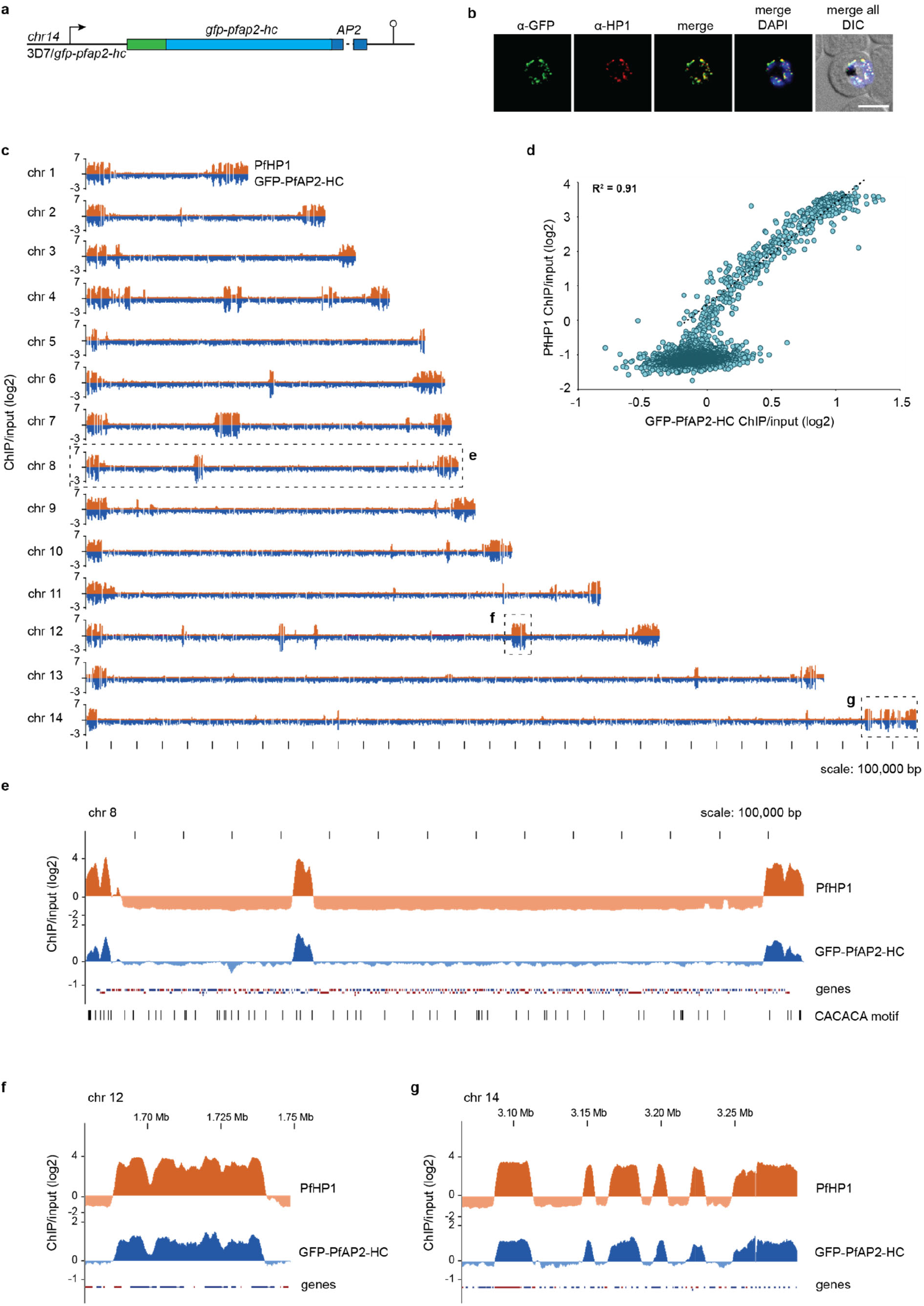
PfAP2-HC associates with PfHP1 throughout the genome. **a** Schematic map of the endogenous *pfap2-hc* locus after introduction of a *gfp* tag by CRISPR/Cas9-mediated gene editing in 3D7/GFP-PfAP2-HC parasites. **b** Representative IFA images of GFP-PfAP2-HC and PfHP1 localisation in 3D7/GFP-PfAP2-HC parasites, 36-44 hpi. Nuclei were stained with DAPI. DIC, differential interference contrast. Scale bar, 5 μm. **c** Log2-transformed α-PfHP1 (orange) and α-GFP (blue) ChIP-over-input ratio tracks obtained from 3D7/GFP-PfAP2-HC schizont stage parasites. α-PfHP1 and α-GFP ChIP tracks have been offset by 2 and 1 respectively to be able to display the full scale of variation. In addition, α-GFP ChIP-seq data is mirrored on a negative scale. Dashed boxes highlight regions that are enlarged in panels e-g. **d**, Scatterplot of average log2-transformed α-PfHP1 and α-GFP ChIP-over-input values for all parasite genes. The depicted regression line is based on heterochromatic genes only (log2 ratio α-PfHP1/input ≥ 0). The coefficient of determination (R^2^) is displayed in the upper left corner. **e** Log2-transformed ChIP-over-input ratio tracks on chromosome 8. The locations of the putative PfAP2-HC binding motif (CACACA) ^35^ are shown below the tracks (FDR < 0.05). Coding sequences are shown as blue (sense strand) and red (antisense strand) boxes. **f**, **g** Log2-transformed ChIP-over-input ratio tracks over sections of chromosome 12 and 14, respectively.

In order to investigate the genome-wide binding profile of GFP-PfAP2-HC and to allow comparison with PfHP1 at high resolution, we performed chromatin immunoprecipitation-sequencing (ChIP-Seq) using α-GFP and α-PfHP1 antibodies to compare binding profiles within the same parasite population. We found that GFP-PfAP2-HC indeed co-localises with PfHP1 throughout the genome (Fig. 1c). To quantify the degree of co-localisation, we computed and compared PfHP1 and PfAP2-HC ChIP-over-input enrichment values in coding regions across the genome (Supplementary Dataset 1). This confirmed a strong correlation (R^2^=0.91) between PfAP2-HC and PfHP1 occupancies across coding regions of all heterochromatic genes (Fig. 1d). In addition, we visualized on all chromosomes the locations of the putative PfAP2-HC target DNA motif (CACACA) as predicted by *in vitro* binding preference of the recombinant PfAP2-HC AP2 DBD ^35^. The CACACA motif showed no enrichment in heterochromatic over euchromatic regions and therefore showed no positional association with the *in vivo* PfAP2-HC binding profile (Fig. 1e). Collectively, these findings show that PfAP2-HC localizes exclusively to PfHP1-defined heterochromatic regions and seems not to bind to the predicted CACACA target motifs *in vivo*.

### PfAP2-HC is not required for heterochromatin maintenance and inheritance

Having shown that PfAP2-HC shares the genome-wide binding profile of PfHP1, we next investigated the function of this ApiAP2 factor by creating a conditional knockdown line employing the FKBP destabilization domain (DD) system. DD-tagged proteins are stabilized in the presence of the small molecule Shield-1 and removal of this ligand leads to protein degradation ^36,37^. We utilized our two-plasmid CRISPR/Cas9 approach to N-terminally tag PfAP2-HC with DDGFP to create the cell line 3D7/DDGFP-PfAP2-HC (Fig. 2a and Supplementary Fig. 2). Limiting dilution cloning resulted in a parasite clone containing the correctly edited locus, which we confirmed by PCR on gDNA (Supplementary Fig. 2). Significant depletion of DDGFP-PfAP2-HC expression in the absence of Shield-1 was verified by live cell fluorescence imaging (Fig. 2b) and Western blot (Fig. 2c and Supplementary Fig. 2). Depletion of DDGFP-PfAP2-HC expression caused no major cell cycle- or proliferation-related phenotypes, nor did it have an effect on sexual conversion rates (Supplementary Fig. 3).

**Figure 2.**
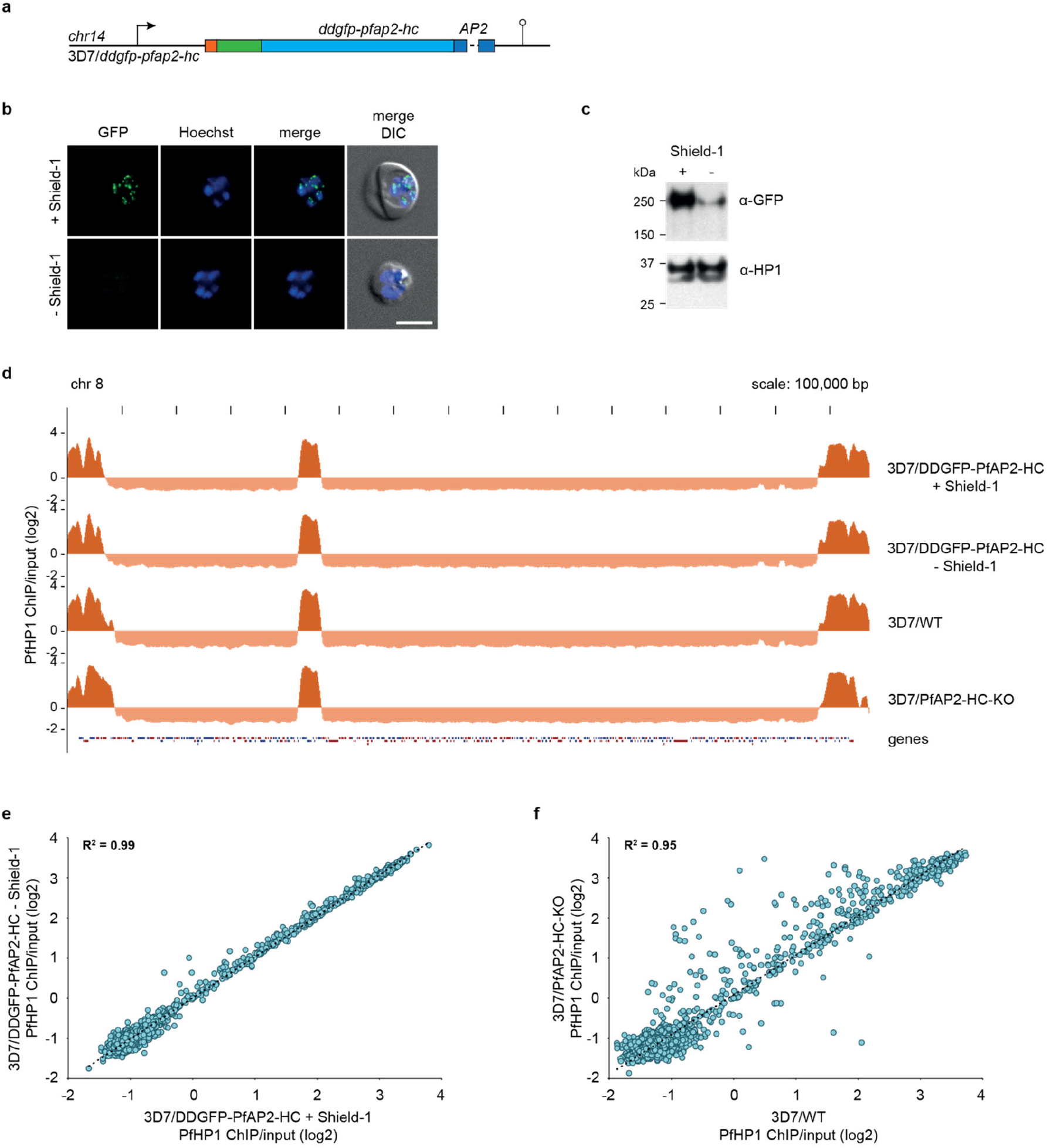
PfAP2-HC depletion does not affect PfHP1 genome-wide coverage. **a** Schematic map of the endogenous *pfap2-hc* locus after CRISPR/Cas9-mediated gene editing to introduce *ddgfp* tag in 3D7/DDGFP-PfAP2-HC parasites. **b** Representative live cell fluorescence images of 3D7/DDGFP-PfAP2-HC schizonts (36-44 hpi) grown in the presence (+) or absence (−) of Shield-1. Nuclei were stained with Hoechst. DIC, differential interference contrast. Scale bar, 5 μm. **c** Western blot showing DDGFP-PfAP2-HC expression levels in 3D7/DDGFP-PfAP2-HC schizonts (36-44 hpi) grown in the presence (+) or absence (−) of Shield-1. PfHP1 expression levels served as a loading control. The full sized blot is available in Supplementary Fig. 2. **d** Log2-transformed α-PfHP1 ChIP-over-input tracks from 3D7/DDGFP-PfAP2-HC schizont stage parasites grown in the presence (+) or absence (−) of Shield-1 (top two tracks). Log2-transformed α-PfHP1 ChIP-over-input tracks from 3D7/WT and 3D7/PfAP2-HC-KO schizonts (bottom two tracks). Coding sequences are shown as blue (sense strand) and red (antisense strand) boxes. **e, f** Scatterplots of average log2-transformed α-PfHP1 ChIP-over-input values at all coding regions in 3D7/DDGFP-PfAP2-HC schizonts grown in the presence (+) or absence (−) of Shield-1 (e) and in 3D7/WT and 3D7/PfAP2-HC-KO schizonts (f). Depicted regression lines are based on heterochromatic genes only (log2 ratio α-PfHP1/input ≥ 0). The coefficient of determination (R^2^) is shown in the upper left corner.

In order to investigate the potential effect of PfAP2-HC depletion on heterochromatin, we grew parasites in the presence or absence of Shield-1 for 13 generations and compared their genome-wide PfHP1 binding profiles by ChIP-Seq. The genome-wide PfHP1 coverage tracks in 3D7/DDGFP-PfAP2-HC parasites grown in the absence or presence of Shield-1 are highly similar (Fig. 2d). Likewise, the genome-wide PfHP1 coverage of coding regions in the two populations is nearly identical (R^2^=0.99, Fig. 2e, Supplementary Dataset 1) showing that depletion of PfAP2-HC has no discernible effect on PfHP1 localization on chromatin. To test whether the lack of obvious loss-of-function phenotypes was due to the residual amounts of DDGFP-PfAP2-HC protein remaining after Shield-1 removal (Fig. 2c), we also generated a PfAP2-HC knockout cell line, 3D7/PfAP2-HC-KO (Supplementary Fig. 4), which we confirmed by PCR on gDNA (Supplementary Fig. 4). 3D7/PfAP2-HC-KO parasites did not show obvious growth-related phenotypic changes either (Supplementary Fig. 4) and maintained PfHP1 occupancy at levels similar to 3D7 wild type (3D7/WT) and 3D7/DDGFP-PfAP2-HC parasites (Fig. 2d). Changes in PfHP1 coverage of some genes were observed in 3D7/PfAP2-HC-KO parasites compared to 3D7/WT and 3D7/DDGFP-PfAP2-HC (Fig. 2f, Supplementary Fig. 4, Supplementary Dataset 1). However, these changes are likely unrelated to the lack of PfAP2-HC expression but rather attributable to clonally variant changes in PfHP1 occupancy as similar differences are observed when comparing different PfAP2-HC-expressing clonal lines (3D7/WT and 3D7/DDGFP-PfAP2-HC) (Supplementary Fig. 4). Together, these results show that PfAP2-HC is required for neither asexual proliferation nor for the maintenance and inheritance of PfHP1-demarcated heterochromatin.

### PfAP2-HC does not act as a transcription factor in blood stage parasites

To identify any possible role of PfAP2-HC in transcriptional regulation we performed a transcriptome-wide microarray time course analysis. We compared 3D7/DDGFP-PfAP2-HC parasites grown in the presence and absence of Shield-1 across five time points throughout the IDC (Fig. 3a). For each of the five time points the paired transcriptome datasets were strongly correlated based on Pearson correlation values, demonstrating highly comparable stage composition across the time course (Fig. 3a and Supplementary Dataset 2). We found no significant difference in gene expression, with no transcripts showing greater than two-fold average fold change in steady-state mRNA abundance between the +Shield-1 and −Shield-1 populations (Fig. 3b), suggesting that PfAP2-HC does not play a dominant role in transcriptional regulation in blood stage parasites and further corroborating the lack of obvious phenotypes associated with PfAP2-HC depletion.

**Figure 3.**
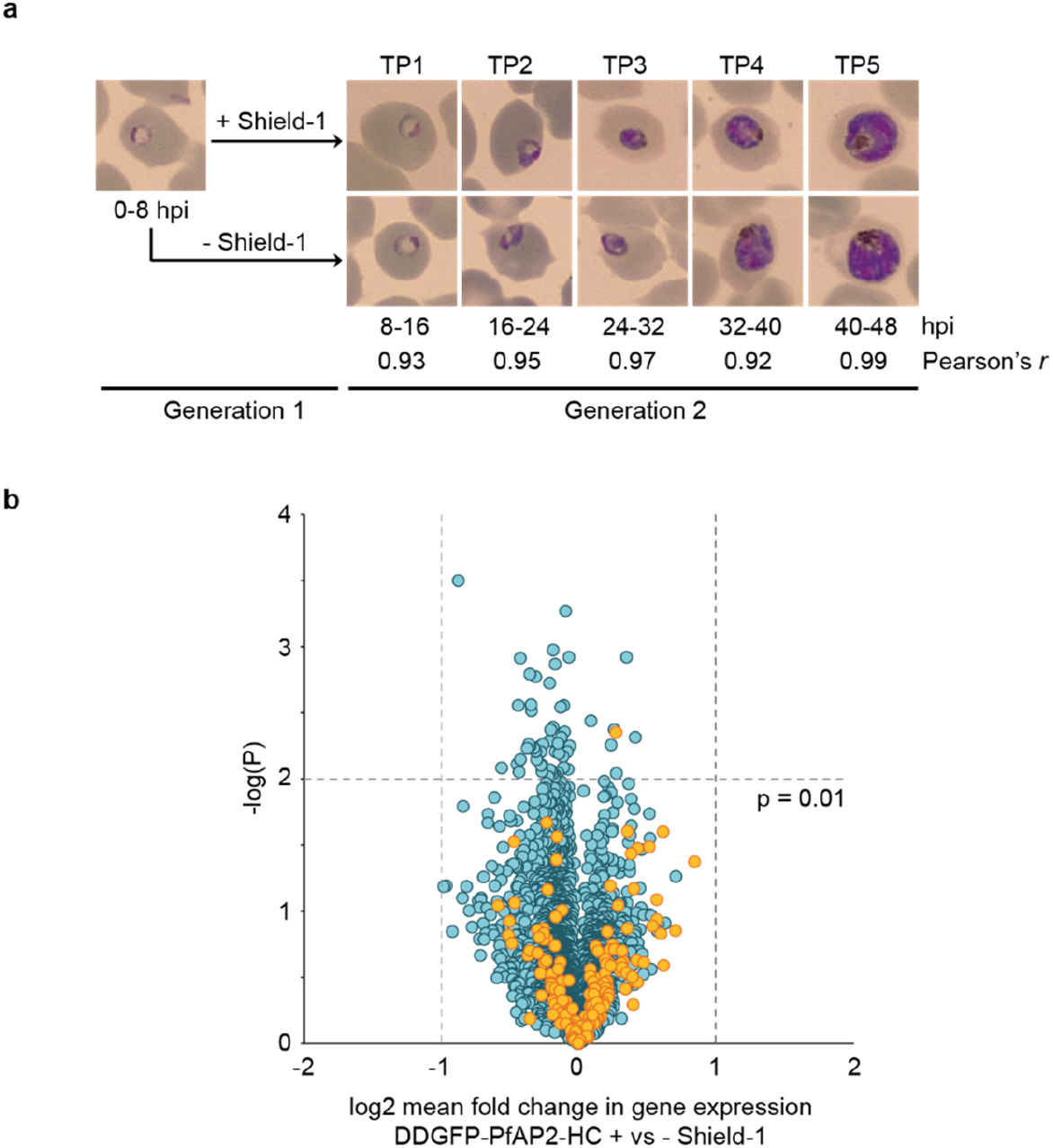
Depletion of PfAP2-HC has no effect on transcription in asexual blood stage parasites. **a** 3D7/DDGFP-PfAP2-HC parasites were grown in the presence (+) or absence (−) of Shield-1 from 0-8 hpi for one generation and sampled for comparative transcriptome analysis at five IDC time points in the subsequent generation. Pearson correlation coefficients *r* indicate the pairwise correlation between the paired transcriptomes of parasites cultured in the presence (+) or absence (−) of Shield-1 for each time point. TP, time point. **b** Volcano plot showing log2 fold changes in relative transcript abundance averaged across the five time points and plotted against significance [−log10(p-value)]. Euchromatic and heterochromatic genes are depicted by blue and orange circles, respectively.

### The AP2 domain of PfAP2-HC is dispensable for targeting PfAP2-HC to heterochromatin

To discern the importance of the single AP2 DBD in targeting PfAP2-HC to heterochromatin we introduced a STOP codon prior to the AP2 domain, replacing amino acid R1319 with a premature STOP codon in 3D7/GFP-PfAP2-HC to create the parasite line 3D7/GFP-PfAP2-HC-ΔDBD (Fig. 4a and Supplementary Fig. 5). PCR on gDNA confirmed successful editing of the locus (Supplementary Fig. 5). The transgenic population consisted of a mixture of parasites with either correctly edited locus or carrying integrated donor plasmid concatemers (Supplementary Fig. 5). Importantly, both recombination events introduce the desired premature STOP codon into the *pfap2-hc* coding sequence. Indeed, Sanger sequencing of the amplified PCR products verified successful introduction of the premature STOP codon in the entire population (Supplementary Fig. 5). The localization of GFP-PfAP2-HC-ΔDBD is comparable to that of GFP-PfAP2-HC by IFA and similarly shares this localization pattern with PfHP1 (Fig. 4b).

**Figure 4.**
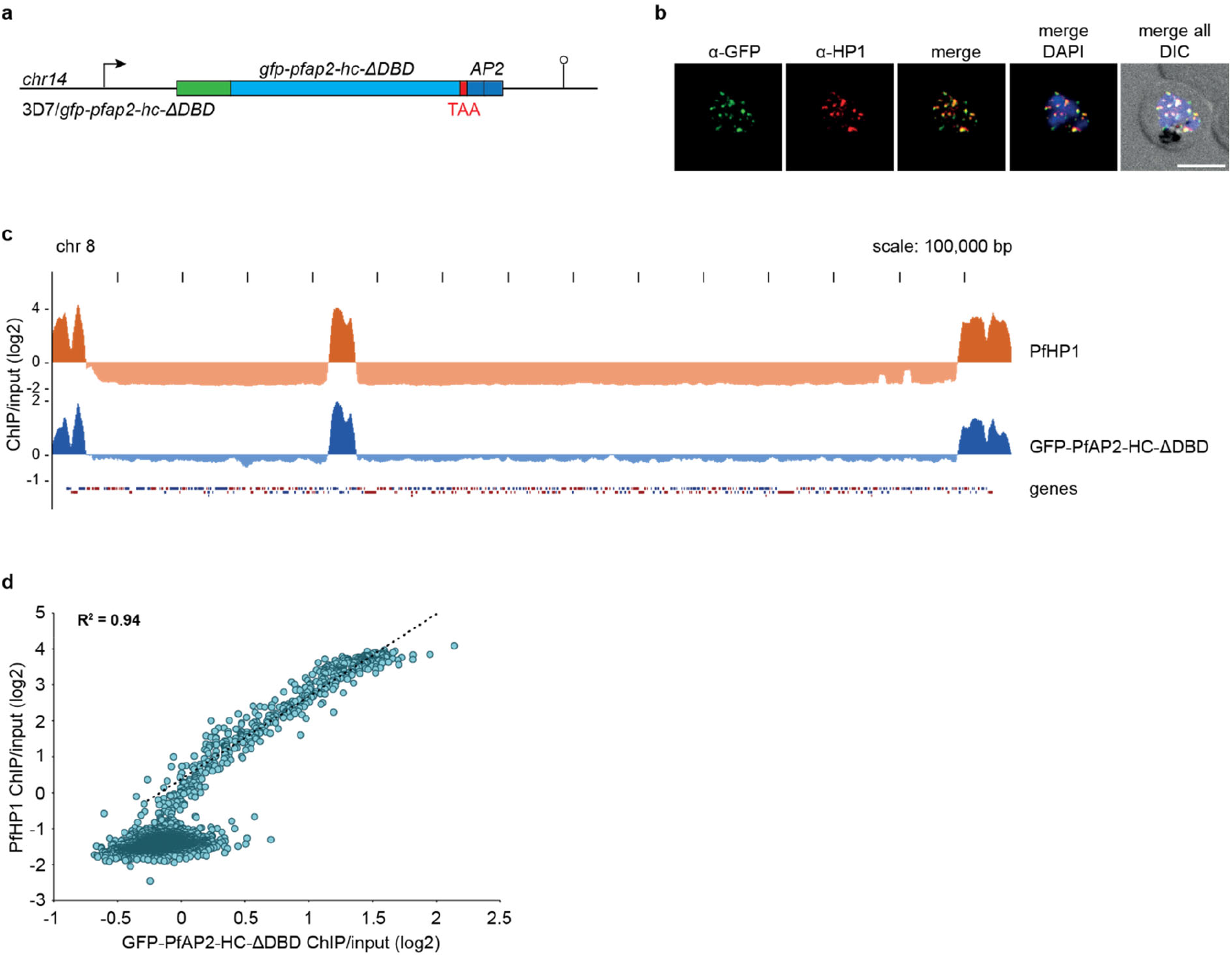
The AP2 domain of PfAP2-HC is not required for heterochromatin targeting. **a** Schematic map of the *gfp-pfap2-hc* locus after CRISPR/Cas9-mediated gene editing to introduce a STOP codon prior to the sequence encoding the AP2 DBD in 3D7/GFP-PfAP2-HC-ΔDBD parasites. **b** Representative IFA images of GFP-PfAP2-HC-ΔDBD and PfHP1 localisation in a developing schizont (36-44 hpi). Nuclei were stained with DAPI. DIC, differential interference contrast. Scale bar, 5 μm. **c** Log2-transformed α-PfHP1 (orange) and α-GFP (blue) ChIP-over-input tracks from 3D7/GFP-PfAP2-HC-ΔDBD schizont stage parasites. **d** Scatterplot of average log2-transformed α-PfHP1 and α-GFP ChIP-over-input values at all coding regions. The regression line is based on heterochromatic genes only (log2 ratio α-PfHP1/input ≥ 0). The coefficient of determination (R^2^) is displayed in the upper left corner.

For a more comprehensive analysis, we again performed ChIP-seq experiments using α-GFP and α-PfHP1 antibodies on 3D7/GFP-PfAP2-HC-ΔDBD parasites. As with full-length GFP-PfAP2-HC, the truncated PfAP2-HC-ΔDBD protein co-localised with PfHP1 throughout the genome with highly correlated enrichment on all heterochromatic genes (Fig. 4c and d, Supplementary Dataset 1) showing that the AP2 DBD of PfAP2-HC is dispensable for its localization to heterochromatin.

### Binding of PfAP2-HC to heterochromatin is PfHP1-dependent

PfAP2-HC is targeted to heterochromatin in the absence of its only recognisable DBD, suggesting a reliance on protein-protein interactions independent of the AP2 domain. To gain insight into this interaction, we tagged PfHP1 with the fluorescent protein mScarlet. In addition, we introduced a sequence encoding the *glms* riboswitch element ^38^ downstream of the STOP codon, such that the resulting *pfhp1-mscarlet* mRNA contains a functional *glms* ribozyme in its 3’ untranslated region. Upon addition of glucosamine (GlcN) to the culture medium, the *glms* ribozyme mediates mRNA cleavage and degradation ^38,39^. We generated this conditional PfHP1 knockdown cassette in the background of the 3D7/GFP-PfAP2-HC clone to create the 3D7/GFP-PfAP2-HC/PfHP1-mScarlet-glmS double transgenic parasite line (Fig. 5a and Supplementary Fig. 6). We confirmed correct editing of the *pfhp1* locus by PCR on gDNA (Supplementary Fig. 6). To investigate the effect of PfHP1 depletion on the localisation of GFP-PfAP2-HC, we split 3D7/GFP-PfAP2-HC/PfHP1-mScarlet-glmS parasites at 0-8 hpi into two populations, adding GlcN to one of them to induce the knockdown of PfHP1-mScarlet expression (+ GlcN), and keeping the other one under stabilising conditions (− GlcN). We performed live cell fluorescence imaging of early schizonts at 32-40 hpi and observed efficient depletion of PfHP1-mScarlet expression in + GlcN conditions (Fig. 5b). Interestingly, upon PfHP1-mScarlet depletion, GFP-PfAP2-HC localised diffusely throughout the nucleoplasm and no longer displayed a punctate perinuclear pattern (Fig. 5b), showing mis-localisation in the absence of PfHP1. The ChIP-seq results presented in Fig. 1 provided no evidence for direct binding of PfAP2-HC to DNA in euchromatic regions. However, this experiment did not allow us to test if PfAP2-HC binds to DNA sequences in heterochromatic regions because its association with PfHP1 would have masked such interactions. Hence, we used the 3D7/GFP-PfAP2-HC/PfHP1-mScarlet-glmS line to ask whether PfAP2-HC binds directly to DNA in the absence of PfHP1. We grew 3D7/GFP-PfAP2-HC/PfHP1-mScarlet-glmS parasites in the presence of GlcN from early ring stages (0-8 hpi) and harvested samples for ChIP-Seq at 40-48 hpi within the same cycle. As expected, we observe a large reduction in PfHP1 enrichment in heterochromatic domains (Fig. 5c). GFP-PfAP2-HC occupancy was massively reduced and in two biologically independent ChIP-seq experiments we could not detect signals over background (Fig. 5c, Supplementary Dataset 1). Together, these results show that PfAP2-HC localisation to heterochromatin is entirely dependent on PfHP1 and no evidence for direct binding of PfAP2-HC to DNA in these regions could be discerned.

**Figure 5.**
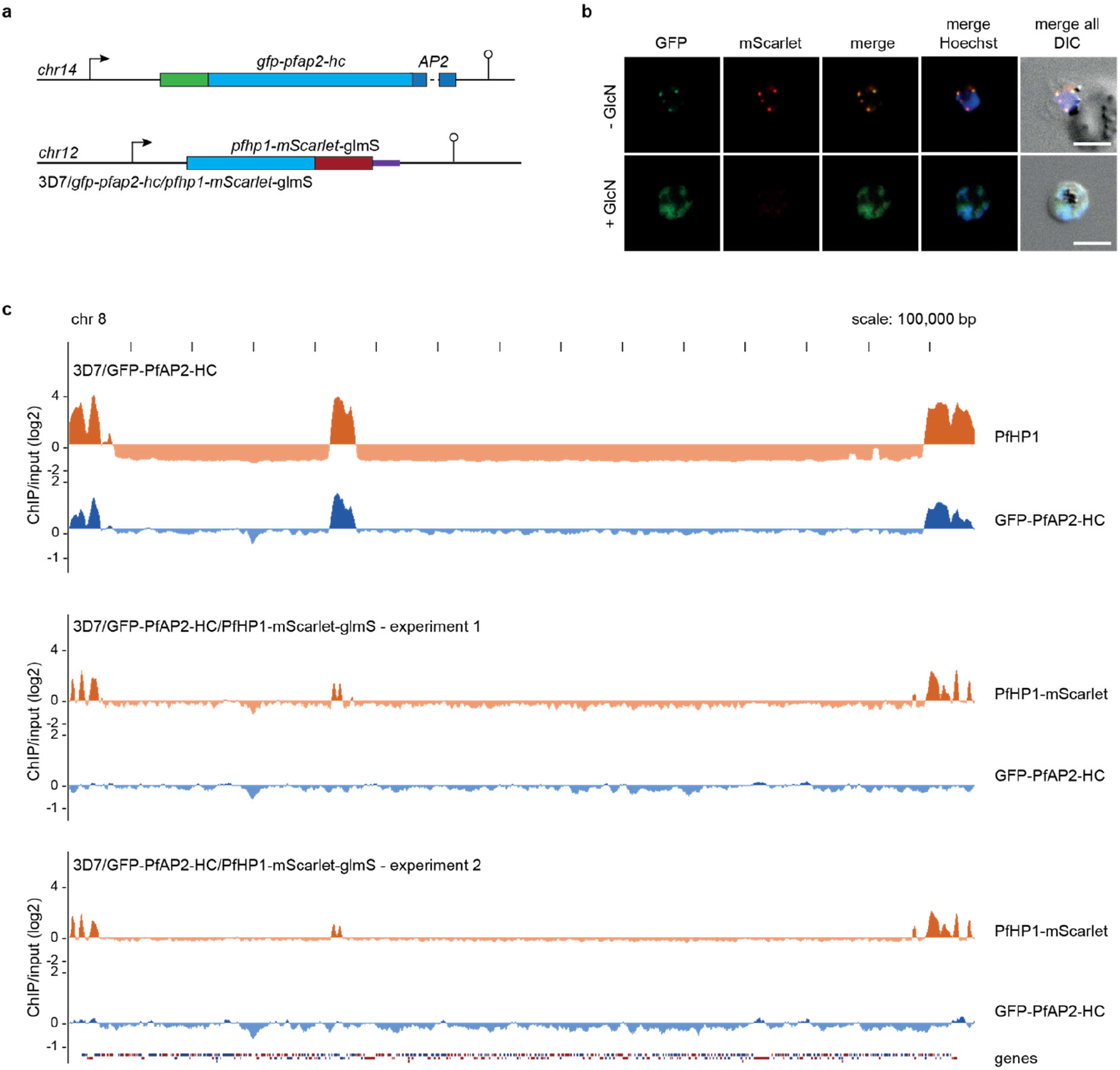
Binding of PfAP2-HC to heterochromatin is PfHP1-dependant. **a** Schematic maps of the endogenous *pfap2-hc* and *pfhp1* loci after CRISPR/Cas9-based editing in 3D7/GFP-PfAP2-HC/PfHP1-mScarlet-glmS parasites. The *pfap2-hc* gene was tagged with *gfp*. The *pfhp1* gene was tagged with the *mScarlet* sequence followed by a *glmS* ribozyme element to allow for detection and conditional expression of PfHP1-mScarlet, respectively. **b** Representative live cell fluorescence images of 3D7/GFP-PfAP2-HC/PfHP1-mScarlet-glmS parasites at 32-40 hpi grown in the absence of GlcN (PfHP1 expressed) or the presence of GlcN (PfHP1 depleted). Nuclei were stained with Hoechst. DIC, differential interference contrast. Scale bar, 5 μm. **c** Log2-transformed α-PfHP1 (orange) and α-GFP (blue) ChIP-over-input tracks from 3D7/GFP-PfAP2-HC schizonts (top, identical to the tracks shown in Fig. 1e) and from 3D7/GFP-PfAP2-HC/PfHP1-mScarlet-glmS schizonts in two independent experiments (middle and bottom).

### PfAP2-HC is likely not involved in heterochromatin formation

We have shown that maintenance and inheritance of heterochromatin was unaffected in both the 3D7/PfAP2-HC-KO null mutant and in the conditional 3D7/DDGFP-AP2-HC loss-of-function mutants after 13 generations of growth under PfAP2-HC-depleted conditions (Fig. 2d). However, factors influencing the initial establishment of heterochromatin can be independent of maintenance and inheritance ^40^. Taking advantage of the fact that conditional knockdown of PfHP1 expression produces progeny consisting of approximately 50% viable heterochromatin-depleted early stage gametocytes and 50% growth-arrested trophozoites ^12^, we investigated whether PfAP2-HC is required for the re-establishment of heterochromatin during gametocyte maturation. To achieve this, we generated a parasite line allowing for the conditional knockdown of both PfHP1 and PfAP2-HC, 3D7/DDGFP-PfAP2-HC/PfHP1-mScarlet-glmS (Fig. 6a and Supplementary Fig. 6). The 3D7/DDGFP-PfAP2-HC/PfHP1-mScarlet-glmS line was obtained by tagging the *pfhp1* gene in the 3D7/DDGFP-AP2-HC clone with mScarlet-glmS as described above (Supplementary Fig. 6). We confirmed correct editing of the *pfhp1* locus by PCR on gDNA (Supplementary Fig. 6). Routine culture of this parasite line in the presence of Shield-1 and absence of GlcN stabilises DDGFP-PfAP2-HC and PfHP1-mScarlet expression, respectively. We divided ring stage parasites into two populations at 0-8 hpi (generation 1), of which one was maintained under stabilising conditions for both proteins, and from the other one Shield-1 was removed to induce DDGFP-PfAP2-HC depletion. At 0-8 hpi in generation 2, we induced the knockdown of PfHP1-mScarlet expression in both populations through addition of GlcN (Fig. 6b). Both populations (DDGFP-AP2-HC stabilised/PfHP1 depleted and DDGFP-AP2-HC depleted/PfHP1 depleted) progressed into generation 3 to produce heterochromatin-depleted sexually committed parasites and growth-arrested trophozoites. On day two of gametocytogenesis, we rescued PfHP1 expression by removal of GlcN from both parasite populations and added 50 mM N-acetyl glucosamine (GlcNac) to prevent multiplication of asexual parasites ^41,42^ that can potentially arise from arrested trophozoites resuming growth after PfHP1 rescue ^12^. We then assessed the re-establishment of perinuclear heterochromatin in presence (+Shield-1) or absence (−Shield-1) of DDGFP-AP2-HC in stage II (64-72 hpi, day 3) (Fig. 6c) and stage V (232-240 hpi, day 10) (Fig. 6d) gametocytes by live cell fluorescence imaging of PfHP1-mScarlet signals. We observed no marked difference in the localisation pattern of PfHP1-mScarlet between gametocytes that express or do not express DDGFP-PfAP2-HC (Fig. 6c and 6d). These observations indicate that PfAP2-HC likely plays no major role in *de novo* heterochromatin formation.

**Figure 6.**
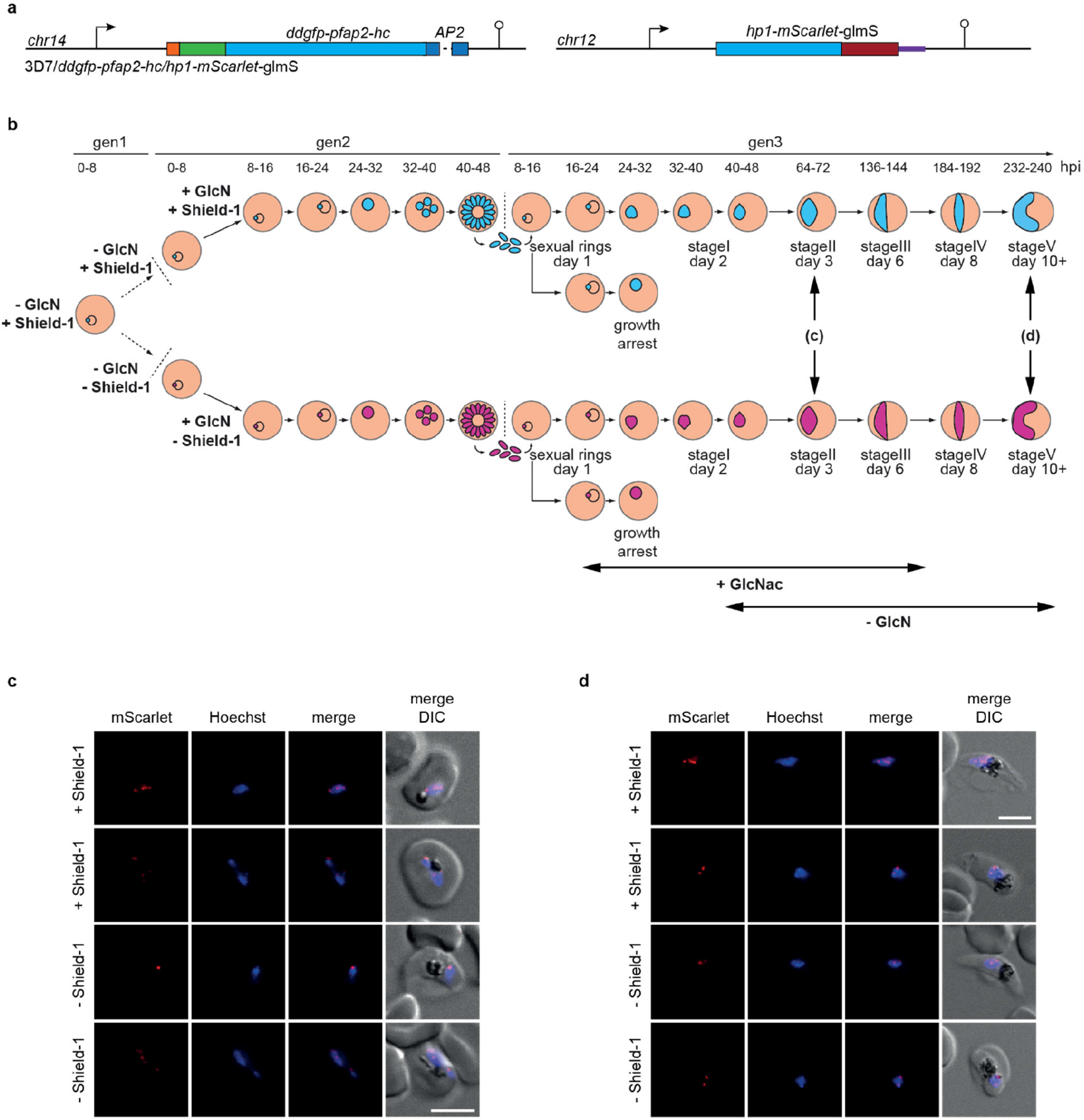
Depletion of PfAP2-HC has no marked effect on re-establishment of heterochromatin. **a** Schematic map of the endogenous *pfap2-hc* and *pfhp1* loci in 3D7/DDGFP-PfAP2-HC/PfHP1-mScarlet-glmS parasites after CRISPR/Cas9-mediated gene editing. The *pfap2-hc* locus was modified to introduce a *ddgfp* tag. The *pfhp1* locus was modified to contain an *mscarlet* tag followed by the *glmS* ribozyme element. **b** Schematic detailing the design of a combined conditional DDGFP-AP2-HC depletion and PfHP1-mScarlet depletion/rescue experiment. Parasites grown in the presence of Shield-1 (+Shield-1) and the absence of glucosamine (−GlcN) exhibit stable expression of both DDGFP-PfAP2-HC and PfHP1-mScarlet. In generation 1, parasites were split into two populations at 0-8 hpi, with Shield-1 removed from one population to induce DDGFP-PfAP2-HC depletion (-Shield-1, magenta parasites) and one population maintained in the presence of Shield-1 (+Shield-1, turquoise parasites). GlcN was added to both populations at 0-8 hpi in generation 2 (+GlcN) to induce PfHP1-mScarlet depletion, which triggers sexual commitment ^12^. In generation 3, 50 mM GlcNAc was added to the ring stage cultures for six days to prevent growth of asexual parasites (depicted with a horizontal arrow) ^41^. Furthermore, GlcN was removed from both populations one day after invasion (i.e. day 2 of gametocytogenesis; stage I gametocytes) (-GlcN, horizontal arrow) to restore PfHP1-mScarlet expression during gametocytogenesis. The double vertical arrows indicate the time points of live cell fluorescence imaging experiments to assess PfHP1-mScarlet localisation in DDGFP-PfAP2-HC-expressing (+Shield-1) and -depleted (−Shield-1) parasites. **c, d** Representative live cell fluorescence images showing PfHP1-mScarlet localisation in stage II gametocytes (c) and stage V gametocytes (d) grown under DDGFP-PfAP2-HC-stabilizing (+Shield-1, upper two panels) and -depleting (−Shield-1, lower two panels) conditions. Nuclei were stained with Hoechst. DIC, differential interference contrast. Scale bar, 5 μm.

## Discussion

Clonally variant gene expression is key to the survival of *P. falciparum* in the human host and is dependent on heterochromatin-mediated gene silencing. PfHP1, as a core component of heterochromatin, is essential for regulating processes as diverse as antigenic variation, invasion pathway switching, commitment to gametocytogenesis and asexual proliferation ^6,12^. Our study characterizes PfAP2-HC, a member of the ApiAP2 family of putative DNA-binding proteins that specifically associates with heterochromatin throughout the genome.

Despite progress towards understanding the heterochromatic landscape of *P. falciparum*, a global view of the dynamic processes occurring to regulate and maintain heterochromatin in this parasite remains elusive. Here, we describe PfAP2-HC as an integral component of heterochromatin, only the second such factor to be characterized after gametocyte development 1 (GDV1) ^33^. GDV1 is not expressed in asexual parasites but only in parasites undergoing sexual commitment. In these cells, GDV1 binds to heterochromatin throughout the genome and destabilizes heterochromatin particularly at the *pfap2-g* locus and early gametocyte markers thus facilitating their expression ^33^. In contrast, PfAP2-HC is expressed and binds to heterochromatin in asexual parasites. Depletion of PfAP2-HC had no effect on PfHP1 localisation suggesting it is not required for heterochromatin maintenance. Factors shown to date to be involved in heterochromatin maintenance in *P. falciparum* consist of histone modifying enzymes, such as the histone deacetylase PfHda2, whose absence leads to the expression of many PfHP1-associated genes including subtelomeric multi-gene families and the internally located *pfap2-g* locus ^43^. The histone deacetylases Sir2A and Sir2B are also required for maintaining *var* gene silencing but do not appear to have a role in regulating *pfap2-g* ^44,45^. The incorporation of PfHP1 into heterochromatin relies on the presence of the histone post-translational modification H3K9me3 ^46^, which is thought to be performed by the histone lysine methyltransferase (HKMT) PfSET3 in *P. falciparum* ^47^. PfSET3 was identified by phylogenetic analysis as a putative orthologue of the SU(VAR)3-9 HKMTs that deposit H3K9me3 marks in model eukaryotes ^47^. PfSET3 was indeed localized to the nuclear periphery in *P. falciparum* ^8,48^, but so far PfSET3 has not been analysed on the functional level and methylation of H3K9 by PfSET3 could not be proven with recombinant protein assays ^47^. In addition to histone modifying enzymes, other putative PfHP1-interacting factors have been identified ^33^, although their role in heterochromatin maintenance is yet to be determined. These include the chromodomain-helicase-DNA-binding protein 1 (PfCHD1), whose homologues are important in chromatin remodeling ^49,50^; both subunits of the FACT histone chaperone, one of which was shown as vital in the production of fertile male gametes in *P. berghei* ^51^; and the Pf14-3-3I reader protein that specifically recognizes phosphorylation of serine 28 on histone 3 ^52^. The manner in which all these chromatin components interact and cooperate to mediate reversible gene silencing in *P. falciparum* is an interesting and equally challenging question for future research.

We also tested whether the absence of PfAP2-HC may influence heterochromatin formation rather than maintenance. Because PfHP1 is essential for the proliferation of asexual parasites, we performed this experiment in gametocytes where PfHP1 is dispensable ^12^. To this end, we first depleted PfHP1 in 3D7/DDGFP-PfAP2-HC/PfHP1-mScarlet-glmS parasites through conditional knockdown of PfHP1 expression, and then rescued PfHP1 expression in the sexual ring stage progeny and visualised the re-establishment of heterochromatic foci in stage II and V gametocytes by fluorescence microscopy based on PfHP1-mScarlet positivity. We did not observe any difference in the localization of PfHP1 between gametocytes expressing or not expressing PfAP2-HC, suggesting that *de novo* formation of heterochromatin occurs independent of PfAP2-HC. This is in keeping with our observation that PfAP2-HC does not seem to bind chromosomal DNA *in vivo* and that the localization of PfAP2-HC is dependent on the presence of PfHP1, as discussed further below. However, we cannot exclude the possibility that a role for PfAP2-HC in nucleating heterochromatin may potentially have been masked by the spreading of heterochromatin from residual PfHP1 that remained bound to chromatin due to incomplete protein knockdown (Fig. 5c).

We showed that the AP2 DBD of PfAP2-HC is not required for correct localization of the protein to heterochromatin. Furthermore, we could not detect direct binding of PfAP2-HC to the predicted CACACA target motifs ^35^ or to other sites in chromosomal DNA by ChIP-Seq, neither in euchromatin nor in heterochromatin, and PfAP2-HC depletion had no effect on gene transcription during the IDC. Together, these results imply that PfAP2-HC does not bind chromosomal DNA *in vivo*, suggesting functional divergence of AP2 domains within the ApiAP2 family. Although DNA-binding motifs were predicted for most AP2 domains *in vitro* ^35^, two of the three AP2 domains of PfAP2-I were recently shown to be dispensable in the IDC and it is unknown if they bind DNA *in vivo* ^21^. It is possible that any direct DNA-binding of PfAP2-HC was below the detection limit of our ChIP-Seq experiments. However, it is perhaps more likely that PfAP2-HC does not bind DNA directly *in vivo*, given its dependence on PfHP1 for correct localization. In fact, because PfAP2-HC interacts with heterochromatin independent of its AP2 domain PfAP2-HC may actually not meant to bind DNA directly; PfAP2-HC would likely recruit heterochromatin to any chromosomal sites it would bind to and thus potentially silence expression of genes that are important for parasite viability. The apparent lack of DNA-binding activity displayed by the PfAP2-HC AP2 domain, and the capacity of PfAP2-HC to localise to heterochromatin in absence of the AP2 domain, suggests that protein-protein interactions involving the large N-terminus of the protein are responsible for targeting PfAP2-HC to heterochromatin. Multiple sequence alignments of AP2-HC orthologues across all human-infecting *Plasmodium* spp. show only 30-36% sequence identity to PfAP2-HC, and this is comparable to the AP2-HC orthologues of rodent-infecting species (31-32%) (Supplementary Fig. 7). High sequence similarity is mainly confined to the AP2 domain itself, which shares ≥ 90% identical amino acids across all species (Supplementary Fig. 7). Interestingly, there is a second semi-conserved region of 172 amino acids within PfAP2-HC with 64-67% sequence identity to the orthologues of other human-infecting species and 53-56% identity to those from rodent-infecting species (Supplementary Fig. 7), which points to an evolutionary conserved feature. One could speculate that this region may be involved in mediating interactions with PfHP1 or other chromatin-associated factors. To date, the role of the non-AP2 region of ApiAP2 proteins has not been explicitly studied. However, given the regulatory roles PfAP2-G ^14,20^, PfAP2-I ^21^ and PfAP2-EXP ^22^ play as transcription factors, as well as the *P. berghei* ApiAP2 factors PbAP2-G ^15^, PbAP2-G2 ^25^, PbAP2-FG ^26^, PbAP2-O ^27,28^, PbAP2-Sp ^29^, PbAP2-L ^30^, it can be assumed that these regions are involved in recruiting transcriptional and epigenetic machinery to the promoters in question. Indeed, co-IP experiments identified the bromodomain protein PfBDP1, PfCHD1 and the FACT complex as potential interaction partners of PfAP2-I ^21^, and truncation of PfAP2-EXP to express only the AP2 domain led to de-regulation of its target genes ^22^. Functional analysis of the semi-conserved region identified in PfAP2-HC may be a promising starting point to begin understanding the role of non-AP2 domain regions in ApiAP2 factor function.

The AP2-HC factor is conserved among all *Plasmodium* spp., which clearly suggests an important role for this factor in the biology of malaria parasites, at least *in vivo*. We obtained a viable PfAP2-HC KO line that lacks any obvious phenotype in asexual blood stage parasites, but we cannot rule out functionally critical roles in other life cycle stages. Indeed, RNA-Seq data shows *pfap2-hc* expression in gametocyte and sporozoite stages (plasmodb.org) ^53–55^. However, the orthologues of PfAP2-HC were successfully disrupted in the rodent malaria parasites *P. berghei* and *P. yoelii*, without discernible growth defects observed during the full life cycle in laboratory animals ^31,32^. These results suggest that functional redundancy or compensatory mechanisms may exist among the ApiAP2 family, as also proposed by Zhang and colleagues ^32^. However, at least in asexual blood stage parasites, we believe mechanisms compensating for loss of PfAP2-HC function are highly unlikely given that the conditional knockdown of PfAP2-HC expression did not result in any transcriptional changes and caused not even temporary defect on parasite growth or multiplication. Beyond this, it is also possible that PfAP2-HC is involved in more subtle processes not studied here, which may not present as immediate phenotypes in loss-of-function mutants but may be crucial for parasite fitness in the field. Examples of such processes are DNA repair/recombination within heterochromatic regions or epigenetic memory/switching frequencies of heterochromatic genes. The heterochromatic subtelomeric regions, which contain several hundred members of multi-copy gene families, recombine at a higher rate than the core genome in *P. falciparum*, resulting in high antigenic diversity within the parasite population ^56–58^. Furthermore, DNA repair mechanisms are generally less efficient in heterochromatin compared to euchromatin and thus contribute to increased mutation rates in these regions ^59,60^. Switches in the transcription of heterochromatic genes creates clonal variation in the expression of surface antigens, invasion factors, nutrient channels or PfAP2-G, allowing the parasite population to adapt to and survive under adverse environmental conditions ^6,61,62^. Activation of silenced heterochromatic genes is linked to local chromatin remodelling, as demonstrated for *var* genes ^12,63,64^, *pfap2-g* ^12,33^ and other clonally variant genes ^65^. As an integral and specific component of heterochromatin, it is at least conceivable that PfAP2-HC may act as a positive or negative regulator of DNA repair or chromatin remodelling processes in heterochromatic regions.

In summary, our study provides a comprehensive analysis of the novel ApiAP2 factor PfAP2-HC, based on the analysis of six different single or double engineered transgenic parasite lines. Along with PfAP2-Tel ^66^ and PfSIP2 ^23^, PfAP2-HC joins the ranks of ApiAP2 factors that do not primarily act as transcriptional regulators. We rather characterised PfAP2-HC as a PfHP1-interacting protein and core component of heterochromatin in *P. falciparum*. We found no evidence for direct binding of PfAP2-HC to chromosomal DNA *in vivo* and show that the localisation of PfAP2-HC to heterochromatin is independent of the AP2 domain but strictly dependent on the presence of PfHP1. While our efforts failed to reveal conclusive insight into PfAP2-HC function, we discovered novel properties of ApiAP2 factors that highlight the functional diversity among the members of this family of putative DNA-binding proteins.

## Materials and Methods

### Parasite culture

*P. falciparum* 3D7 parasites were cultured as described ^67^ in RPMI Medium 1640 [+] L-Glutamine (Life Technologies) supplemented with 25 mM HEPES, pH 6.72, 100 mM hypoxanthine, 24 mM sodium bicarbonate and 0.5% Albumax II. 2 mM choline chloride was added to the medium to reduce sexual commitment rates ^68^. Synchronisation of parasite growth was achieved by repeated sorbitol treatments of ring stage parasites ^69^. Parasite cultures were kept at 37 °C under a gaseous mixture of 4% CO_2_, 3% O_2_ and 93% N_2_.

### Transfection constructs

Transgenic cell lines were generated by CRISPR/Cas9-based genome editing using a set of plasmids recently described ^33^. All sgRNA target sequences were identified using CHOPCHOP ^70–72^. 3D7/GFP-PfAP2-HC parasites were created using a two-plasmid approach, consisting of a CRISPR/Cas9 transfection vector pH_gC-*ap2-hc-5'*-1 and the donor plasmid pD_*gfp-pfap2-hc*. The pH_gC-*ap2-hc-5'*-1 plasmid was created by annealing complementary oligonucleotides (sgRNA_ap2-hc-5'-1_F and sgRNA_ap2-hc-5'-1_R) encoding the sgRNA target sequence sgt_ap2-hc-5’-1 (gaaacacataacgagcttaa; positioned at bps +124 to +143 of the *pfap2-hc* coding sequence) and ligating them into the *Bsa*I-digested pH-gC plasmid ^33^. The pD_*gfp-pfap2-hc* donor plasmid was produced by Gibson assembly ^73,74^ of four PCR products encoding (1) the plasmid backbone amplified from pUC19 using primers PCRA_F and PCRA_R ^33^, (2) a 5’ homology region (HR) spanning 575 bp of the *pfap2-hc* upstream region amplified from 3D7 gDNA using primers ap2-hc-5'_HR1_F and ap2-hc-5'_HR1_R, (3) the *gfp* coding sequence amplified from plasmid pD_*ap2g-gfp-dd-glmS* ^33^ using primers ap2-hc-5'_GFP_F and ap2-hc-5'_GFP_R, and (4) a 745 bp 3’ HR corresponding to the *pfap2-hc* coding region +3 to +748. Fragment 4 was ordered as synthetic sequence (GenScript) with the first 274 bp recodonised and was amplified from plasmid pUC57-re-*ap2-hc*-1 using primers ap2-hc-5'_HR2_re_F and ap2-hc-5'_HR2_re_R.

The 3D7/DDGFP-PfAP2-HC parasite line was generated using plasmids pH_gC-*ap2-hc-5'*-2 and pFDon_*ddgfp-pfap2-hc*. To generate pH_gC-*ap2-hc-5'*-2, complementary oligonucleotides (sgRNA_ap2-hc-5'-2_F and sgRNA_ap2-hc-5'-2_R) encoding the sgRNA target sequence sgt_ap2-hc-5’-2 (taacaatatttctgtatcta; positioned at bps +36 to +55 of the *pfap2-hc* coding sequence) were annealed and ligated into the *Bsa*I-digested pH-gC plasmid ^33^. pFDon_*ddgfp-pfap2-hc* donor plasmid was created by Gibson assembly of four fragments: (1) the pFDon plasmid ^33^ digested with *Hind*III and *Eco*RI, (2, 3) the 5’ HR and the *gfp*-3’HR were amplified from plasmid pD_*gfp-pfap2-hc* (described above) with primers ap2-hc-5'_HR3_F and ap2-hc-5'_HR3_R, and ap2-hc-5'_HR4_F and ap2-hc-5'_HR4_R, respectively. The final fragment 4 encoding the FKBP destabilizing domain (*dd*) sequence (plus C-terminal TGSS linker) was amplified from pD_*ap2g-gfp-dd-glmS* ^33^ using primers ap2-hc-5'_DD_F and ap2-hc-5'_DD_R.

Parasite line 3D7/GFP-PfAP2-HC-ΔDBD was created by CRISPR/Cas9 editing of 3D7/GFP-PfAP2-HC parasites using plasmids pH_gC-*ap2-hc-3'* and pD_*gfp-pfap2-hc-*Δ*DBD*. The sgRNA-encoding oligonucleotides sgRNA_ap2-hc-3'_F and sgRNA_ap2-hc-3'_R were annealed and ligated into the *Bsa*I-digested pH-gC plasmid ^33^, as above, to create pH_gC-*ap2-hc-3'*. The sgRNA target sequence sgt_ap2-hc-3’ (ctagacaaaaggctattgaa) is positioned at bps +4070 to +4089 of the *pfap2-hc* coding sequence. To create pD_*gfp-pfap2-hc-*Δ*DBD*, a synthetic DNA sequence (GenScript), corresponding to the *pfap2-hc* coding sequence +3528 to +4125 with the intron removed and the sequence +3954 to +4125 recodonised (plasmid pUC57-re-*ap2-hc*-2). To introduce a STOP codon prior to the sequence encoding the AP2 DBD two overlapping PCR fragments (1, 2) were amplified from pUC57-re-*ap2-hc*-2 using primers ap2-hc-3'_HR1_re_F and ap2-hc-3'_HR1_re_R, and ap2-hc-3'_HR2_re_F and ap2-hc-3'_HR2_re_R, respectively. The ap2-hc-3'_HR1_re_R and ap2-hc-3'_HR2_re_F primers introduce a TAA STOP codon at amino acid position 1319 (R1319*). Fragments (1, 2) were assembled together with fragment 3 representing the plasmid backbone amplified from pUC19 using primers PCRA_F and PCRA_R ^33^, and fragment 4 representing the 3’ HR beginning at nucleotide +4088 of the *pfap2-hc* coding sequence and ending 852 bp downstream of the native STOP codon (76 bp into the neighbouring gene PF3D7_1456100) and amplified from 3D7 gDNA using primers ap2-hc-3'_HR3_F and ap2-hc-3'_HR3_R.

3D7/GFP-PfAP2-HC/PfHP1-mScarlet-glmS and 3D7/DDGFP-PfAP2-HC/PfHP1-mScarlet-glmS parasite lines were generated by editing the endogenous *pfhp1* locus in parasites lines 3D7/GFP-PfAP2-HC and 3D7/DDGFP-PfAP2-HC, respectively. The recently published CRISPR/Cas9 plasmid pBF-gC-guide250 ^75^ was used in combination with the donor plasmid pD_*hp1-mScarlet-glmS*. The donor construct was created by joining five fragments in a Gibson assembly consisting of (1, 2) previously described 5’ and 3’ HRs amplified from plasmid pD-PfHP1-KO ^75^, using primers F158 and hp1_HR1_R, and hp1_HR2_F and R163, respectively. Fragment 3, consisting of a *P. falciparum* codon-optimised *mScarlet* sequence with an N-terminal GSAG linker, was amplified from the plasmid pD_*ap2-g-mScarlet* (Brancucci et al., manuscript in preparation) using primers hp1_mScarlet_F and hp1_mScarlet_R. The *glmS* sequence (fragment 4) was amplified from plasmid pL6-3HA_glmS-246 (kind gift from Dave Richard) using primers hp1_glmS_F and hp1_glmS_R, and finally, the plasmid backbone (fragment 5), was amplified from pUC19 with primers PCRA_F and PCRA_R ^33^.

The 3D7/PfAP2-HC-KO cell line was created using a single plasmid CRISPR/Cas9 approach. The mother plasmid p_gC ^33^ formed the backbone to create p_gCH-*pfap2-hc*-KO. p_gC was digested with *Bam*HI and *Hind*III and used in a Gibson assembly with (1) a 5’ HR spanning bps +128 to +533 of the *pfap2-hc* coding sequence, amplified from 3D7 gDNA with primers ap2-hc-KO_HR1_F and ap2-hc-KO_HR1_R, (2) a *hdhfr* expression cassette amplified from plasmid p_gCH-*gdv1*-asKO ^33^ (primers ap2-hc-KO_hDHFR_F and ap2-hc-KO_hDHFR_R), and (3) a 3’ HR spanning bps +3042 to +3460 of the *pfap2-hc* coding sequence, amplified from 3D7 gDNA using primers ap2-hc-KO_HR2_F and ap2-hc-KO_HR2_R. The resulting plasmid, p_gCH-*pfap2-hc*-KO-pre, was digested with *Bsa*I and the sgRNA-encoding sequence sgt_ap2-hc-KO (cgttgtactagtaacattgg; position +1724 to +1743 of the *pfap2-hc* coding sequence, negative strand) was created by annealing the complementary oligonucleotides sgRNA_ap2-hc-KO_F and sgRNA_ap2-hc-KO_R, and ligated into the *Bsa*I site creating the final p_gCH-*pfap2-hc*-KO plasmid.

Oligonucleotide sequences used in cloning are provided in Supplementary Table S1.

### Transfection and transgenic cell lines

*P. falciparum* parasite transfections were carried out as described ^33,76^. A total of 100 μg plasmid DNA (two-plasmid CRISPR/Cas9 approach: 50 μg of each plasmid; single-plasmid CRISPR/Cas9 approach: 100 μg plasmid) was transfected into 3D7/WT or previously engineered parasites and the cultures allowed to recover for 24 hours by growth in drug-free culture medium. Selection of transgenics was then initiated by the addition of 4 nM WR99210 for a total of six days for pH-derived plasmids, or continuously for plasmid p_gCH-*pfap2-hc*-KO. Parasites transfected with the pBF-gC-guide250 construct were selected with 2.5 μg/mL blasticidin-S-hydrochloride for a total of ten days. 3D7/DDGFP-PfAP2-HC parasites were cultured in the presence of 700 nM Shield-1 (+Shield-1) unless otherwise stated. Induction of PfHP1 depletion in parasite lines 3D7/GFP-PfAP2-HC/PfHP1-mScarlet-glmS and 3D7/DDGFP-PfAP2-HC/PfHP1-mScarlet-glmS was achieved by the addition of 2.5 mM glucosamine (GlcN, Sigma #G4875). Limiting dilution cloning was carried out on parasites lines 3D7/GFP-PfAP2-HC, 3D7/DDGFP-PfAP2-HC and 3D7/GFP-PfAP2-HC-ΔDBD as described ^77^. Successful gene editing was confirmed by PCR on gDNA using primers listed in Supplementary Table S2.

### Fluorescence microscopy

Live cell fluorescence imaging was performed as previously described ^78^ with the minor modification of nuclear staining with Hoechst (Merck) instead of DAPI at a final concentration of 5 μg/ml. IFAs were carried out on methanol-fixed cells using primary antibodies mouse mAb α-GFP (Roche Diagnostics #11814460001), 1:100 and rabbit α-PfHP1 ^12^, 1:100. Secondary antibodies Alexa Fluor 488-conjugated α-mouse IgG (Invitrogen #A11001) and Alexa Fluor 568-conjugated α-rabbit IgG (Invitrogen #A11011) were used, each at 1:250 dilution. Nuclei were stained during slide preparation with Vectashield containing DAPI (Vector Laboratories). Images were acquired on a Leica DM 5000B microscope with a Leica DFC 345 FX camera using the Leica application suite (LAS) software. Image processing was carried out using Fiji ^79^. For each experiment, all images were acquired and processed with identical settings.

### Western blot

Whole parasite protein extracts were prepared by first releasing parasites from the iRBC by saponin lysis (0.15% in PBS) followed by suspension of the parasite pellet in UREA/SDS lysis buffer [(8 M Urea, 5% SDS, 50 mM Bis-Tris, 2 mM EDTA, 25 mM HCl, pH 6.5, 1 mM DTT, 1x protease inhibitor (Roche)] and separated on a NuPage 3-8% Tris-Acetate gel (Novex) using NuPage MES SDS Running Buffer (Novex). Proteins were detected with primary antibodies mouse mAb α-GFP (Roche Diagnostics #11814460001), 1:1000 and rabbit α-PfHP1 ^12^, 1:5000, and secondary antibodies α-mouse IgG (H&L)-HRP (GE healthcare #NXA931), 1:5000 and α-rabbit IgG (H&L)-HRP (GE Healthcare #NA934), 1:5000. Chemiluminescence signal was detected using KPL LumiGLO Reserve Chemiluminescent Substrate Kit (SeraCare #5430-0049).

### Chromatin immunoprecipitations

Parasite cultures were synchronised to obtain an eight-hour growth window and harvested at peak PfAP2-HC expression at 36-44 hpi ^34^ from a 30 ml culture at 5% haematocrit and 4-5% parasitemia. Parasites were crosslinked with 1% formaldehyde for 10 min at 37 °C before quenching with 0.125 M glycine. The RBC membrane was lysed with 0.15% saponin and cytoplasmic lysis buffer [(CLB: 20 mM Hepes, 10 mM KCl, 1 mM EDTA, 1 mM EGTA, 0.65% NP-40, 1 mM DTT, 1x protease inhibitor (Roche)] was added to the parasite pellet to isolate nuclei. Nuclei were then snap-frozen in liquid nitrogen in CLB supplemented with 50% glycerol and stored at −80 °C. Chromatin isolation, shearing and immunoprecipitation was performed according to previously published protocols ^33^. To prepare chromatin, frozen nuclei were thawed, pelleted and resuspended in 150 μl sonication buffer [(50 mM Tris pH 8, 1% SDS, 10 mM EDTA, 1x protease inhibitor (Roche)], and were sonicated for 20 cycles of 30 sec ON/30 sec OFF (setting high, BioruptorTM Next Gen, Diagenode) to shear DNA to fragments of 100-600 bps. Fragment size was confirmed by de-crosslinking a 15 μl aliquot and visualising the purified DNA on a 2% agarose gel. ChIPs were performed by combining sonicated chromatin (500 ng DNA content) with either 1 μg mouse mAb α-GFP (Roche Diagnostics #11814460001) or 1 μg rabbit α-PfHP1 ^12^ in incubation buffer [(5% Triton-X-100, 750 mM NaCl, 5 mM EDTA, 2.5 mM EGTA, 100 mM Hepes pH 7.4, 0.2% bovine serum albumin, 1x protease inhibitor (Roche)] containing 10 μl protA and 10 μl protG Dynabeads (Life Technologies, #10008D and #10009D, respectively) in a total reaction volume of 300 μl. ChIP samples were incubated overnight at 4 °C with rotation. Beads were washed for 5 min at 4 °C, with rotation, with 400 μl wash buffers as follows: 2x wash buffer 1 (0.1% SDS, 0.1% DOC, 1% Triton-X100, 150 mM NaCl, 1 mM EDTA, 0.5 mM EGTA, 20 mM Hepes pH 7.4), 1x wash buffer 2 (0.1% SDS, 0.1% DOC, 1% Triton-X100, 500 mM NaCl, 1 mM EDTA, 0.5 mM EGTA, 20 mM Hepes pH 7.4), 1x wash buffer 3 (250 mM LiCl, 0.5% DOC, 0.5% NP-40, 1 mM EDTA, 0.5 mM EGTA, 20 mM Hepes pH 7.4), 2x wash buffer 4 (1 mM EDTA, 0.5 mM EGTA, 20 mM Hepes pH 7.4). Immunoprecipitated chromatin was eluted from the beads by shaking at room temperature for 20 min in 200 μl elution buffer (1% SDS, 0.1 M NaHCO_3_) and de-crosslinked at 45 °C overnight in 1% SDS, 0.1 M NaHCO_3_ and 1 M NaCl. Simultaneously, 30 μl of sonicated input chromatin was de-crosslinked under the same conditions. DNA was purified with QIAquick MinElute PCR columns (Qiagen). For each ChIP-seq experiment, twenty separate α-GFP ChIPs or four separate α-PfHP1 ChIPs were combined, with the exception of 3D7/GFP-PfAP2-HC/PfHP1-mScarlet-glmS α-PfHP1 ChIP-seq for which eight separate ChIPs were combined.

### High throughput sequencing and data analysis

The obtained ChIPed DNA fragments were used to generate Illumina sequencing libraries according to Filarsky, et al. ^33^. In brief, 1 ng of α-PfHP1 ChIP, α-GFP ChIP, or input DNA were end-repaired with T4 DNA polymerase (NEB, M0203L), Klenow DNA polymerase (NEB, M0210L), and T4 Polynucleotide Kinase (NEB, M0201L). The 3’ends of end-repaired DNA were extended with an A-overhang with 3’ to 5’ exonuclease-deficient Klenow DNA polymerase (NEB, M0212L). The resulting fragments were ligated to Nextflex 6bp adaptors (Bio Scientific, #514122) with the use of T4 DNA ligase (Promega, M1804). The libraries were amplified using an AT-rich optimized KAPA protocol using KAPA HiFi HotStart ready mix (KAPA Biosystems, KM2602), NextFlex primer mix (Bio Scientific, #514122) with the following PCR program: 98°C for 2 min; four cycles of 98°C for 20 sec, 62°C for 3 min; 62°C for 5 min. The fragments originating from mono-nucleosomes + 125 bp NextFlex adapter were selected using 2% E-Gel Size Select agarose gels (Invitrogen, #G6610-02) and amplified by PCR for nine cycles using the above conditions. Libraries were purified and adapter dimers removed with Agencourt AMPure XP beads purification using a 1:1 library:beads ratio (Beckman Coulter, #A63880). ChIP-seq libraries were sequenced on the Illumina NextSeq 500 system with a 20% phiX spike-in (Illumina, FC-110-3001) to generate 75 bp single-end reads (NextSeq 500/550 High Output v2 kit). The quality of the resulting reads were checked with FastQC (V0.11.8) and the reads were mapped against the *P. falciparum* 3D7 reference genome from PlasmoDB v26 (www.plasmodb.org) using BWA samse (v0.7.17-r1188) ^80,81^. Mapped reads originating from the mitochondrial and apicoplast genome, multi-mapping reads, and reads having a mapping quality below 15 were removed (SAMtools v1.9) ^82^ leaving between 4.8 and 25.4 million reads (note that replicate 2 of 3D7/GFP-PfAP2-HC/PfHP1-mScarlet-glmS cultured in presence of GlcN (Fig. 5d) is based on 2.0 million reads). Before visualising the ChIP-seq data in the UCSC Genome browser (Fig. 2d, 4d, 5d) or Signalmap (version 2.0.0.5) (Fig. 1d), the libraries were normalised to the total amount of reads, reads per million, with bedtools genomeCoverageBed (v2.27.1) and the enrichment over the input sample was calculated by dividing ChIP sample with input sample read counts, which subsequently was log2 transformed (one pseudo count was added to avoid division by zero) ^83,84^. Within the UCSC genome browser tracks were smoothened (8) and the windowing function was set as ‘mean’. To enable the visualisation of all chromosomes together Signalmap (version 2.0.0.5) was used. The average log2 Chip-over-input value was calculated in sequential 1kb windows to smoothen and compress the data set. PfHP1 values are visualised on the positive scale shifted up by 2 and PfAP2-HC (GFP) values are visualised in the negative scale shifted up by 1. The enrichment tracks were shifted to show the complete tracks. To show the genome-wide colocalization of PfHP1 and PfAP2-HC occupancy and comparison between cell lines the average log2 Chip-over-input ratios at coding genes were calculated using bedtools genomeCoverageBed (v2.27.1) ^84^ (Supplementary Dataset 1) and visualised with Excel 2016. The position of putative AP2-HC binding sites (i.e. CACACA motifs) has been defined using the position weight matrix of the CACACA motif as described by Campbell, et al. ^35^ and searching the *P falciparum* genome for matching sequences (fdr<0.05) with the use of the gimme scan tool of GimmeMotifs ^85^.

### Flow cytometry

Synchronous 3D7/DDGFP-PfAP2-HC and 3D7/WT parasites were split at 0-8 hpi to 0.1% parasitaemia and cultured either in the presence of 700 nM Shield-1 (+ Shield-1) or absence of Shield-1 (− Shield-1) during the duration of the multiplication assay. Synchronous 3D7/PfAP2-HC-KO parasites at 0-8 hpi were diluted to 0.1% parasiteamia. After 24 hours (24-32 hpi) parasite DNA was stained with SYBR Green DNA stain (1: 10,000) (Invitrogen #S7563) for 30 min at 37 °C and the fluorescence intensity was measured using a MACS Quant Analyzer 10 (at least 200,000 RBCs were measured per sample) to determine the parasitaemia (day 1). Measurements were repeated on day 3 and day 5 at 24-32 hpi. Data were analysed using the FlowJo_v10.6.1 software. Gating was performed to remove debris smaller than cell size, to include only single measurement events and to separate uninfected from infected RBCs based on the SYBR Green intensity of an uninfected RBC control sample (the gates for ‘cells’, ‘singlets’ and ‘parasites’, respectively, are shown in Supplementary Fig. 2).

### Quantification of sexual conversion rates

3D7/DDGFP-PfAP2-HC and 3D7/WT parasites were synchronised with sorbitol to a 6-hour time window and again in the next generation after RBC invasion at 0-6 hpi, after which each culture was split and one half was maintained in the presence of Shield-1 (+ Shield-1) and the other half was cultured in the absence of Shield-1 (− Shield-1). Eighteen hours later (18-24 hpi), cultures were split again (1-2% parasitaemia, 2.5% haematocrit) and grown in either minimal fatty acid (mFA) medium [RPMI Medium 1640 [+] L-Glutamine (Life Technologies) supplemented with 25 mM HEPES, pH 6.72, 100 mM hypoxanthine, 24 mM sodium bicarbonate, 0.39% fatty acid-free BSA (Sigma #A6003), 30 μM oleic acid and 30 μM palmitic acid], which induces sexual commitment, or mFA medium supplemented with 2 mM choline chloride (mFA+CC), which inhibits sexual commitment ^68^. After 24 hours (42-48 hpi), mFA or mFA+CC medium was replaced with the standard Albumax medium used throughout this study (see above). After invasion into new RBCs, 50 mM N-acetylglucosamine (GlcNAc) was added to the cultures at 24-30 hpi (day 1 of gametocytogenesis) to prevent asexual parasite multiplication ^41,42^. The resulting gametocyte cultures were maintained in the presence of GlcNAc for a further 3 days. Parasitaemia on day 1 of gametocytogenesis (mixture of asexuals and sexual ring stage parasites) and gametocytemia on day 4 of gametocytogenesis were counted and sexual conversion rates were calculated (gametocytaemia [%] / parasitaemia [%] * 100 = sexual conversion rate [%]).

### Microarray Experiments and Data Analysis

3D7/DDGFP-PfAP2-HC parasites, continuously cultured in the presence of Shield-1, were synchronised with sorbitol to obtain an eight-hour growth window (16-24 hpi) and again in the next generation after RBC invasion at 0-8 hpi. The culture was then split at 8-16 hpi and one half was maintained in the presence of Shield-1 (+Shield-1) and the other half was cultured in the absence of Shield-1 (-Shield-1) to achieve DDGFP-PfAP2-HC depletion. The parasites proceeded through the IDC and were harvested for total RNA extraction in the subsequent generation at the following time points: TP1 (8-16 hpi), TP2 (16-24 hpi), TP3 (24-32 hpi), TP4 (32-40 hpi) and TP5 (40-48 hpi). RNA isolation and cDNA synthesis were performed as previously described ^86^. Cy5-labelled sample cDNAs were hybridised against a Cy3-labelled cDNA reference pool prepared from 3D7 wild-type parasites ^12^. Equal amounts of Cy5- and Cy3-labelled samples were hybridised on a *P. falciparum* 8×15K Agilent gene expression microarray (GEO platform ID GPL15130)^87^ for 16 hours at 65°C in an Agilent hybridisation oven (G2545A). Slides were scanned using the GenePix scanner 4000B and GenePix pro 6.0 software (Molecular Devices). The raw microarray data representing relative steady state mRNA abundance ratios between each test sample and the reference pool (Cy5/Cy3 log2 ratios) were subjected to lowess normalization and background filtering as implemented by the Acuity 4.0 program (Molecular Devices). Flagged features and features with either Cy3 or Cy5 intensities lower than two-fold the background were removed. Log2 ratios for multiple probes per gene were averaged and genes recognized by non-uniquely mapping probes were removed from the dataset. Transcripts showing expression values in at least four of the five samples harvested for each time course were included for downstream analysis to identify genes differentially expressed [mean fold change cut-off > 2; p-value cut-off 0.01 (paired two-tailed Student’s t-test)] between control (+Shield-1) and DDGFP-AP2-HC-depleted (-Shield-1) parasites. The processed microarray dataset is listed in Supplementary Dataset 2.

### Induction of gametocytogenesis by conditional depletion of PfHP1

3D7/DDGFP-PfAP2-HC/PfHP1-mScarlet-glmS parasites were synchronised to obtain an eight-hour growth window and re-synchronised in the following cycle at 0-8 hpi (generation 1). The culture was split into two populations, one grown in the presence of Shield-1 and one in the absence of Shield-1. Both parasite populations were synchronised again at 0-8 hpi in the following cycle (generation 2) and 2.5 mM glucosamine (GlcN, Sigma #G4875) was added to induce PfHP1 depletion (+Shield-1/+GlcN and −Shield-1/+GlcN). Parasites progressed through generation 2 and in generation 3, GlcN was removed from both populations and parasites were cultured on serum medium (0.5% Albumax replaced with 10% human serum) containing 50 mM N-acetylglucosamine (GlcNAc) to prevent asexual parasite multiplication ^41,42^. Live cell fluorescence imaging was performed in stage II and stage V gametocytes to observe PfHP1-mScarlet signals between +Shield-1 (DDGFP-AP2-HC-expressing) and −Shield-1 (DDGFP-AP2-HC-depleted) parasites. The experimental design is depicted schematically in Fig. 6b.

### Multiple sequence alignment of AP2-HC orthologues

Orthologues of PfAP2-HC (PF3D7_1456000) were identified on www.plasmodb.org ^53^ from *P. vivax* (PVX_117665), *P. knowlesi* (PKNH_1225800), *P. malariae* (PmUG01_12060900), *P. ovale curtisi* (PocGH01_12058800), *P. berghei* (PBANKA_1319700), *P. yoelii* (PY17X_1323500) and *P. chabaudi* (PCHAS_1323000) and their full length amino acid sequences were aligned using Clustal X2.1 ^88^ multiple sequence alignment on default settings. A tree was generated using Clustal X2.1 ^88^ on default settings and the resulting identity matrix was tabulated. The AP2 domains of each orthologue were aligned separately and an identity matrix generated. A semi-conserved domain (spanning amino acids 995-1126 of PfAP2-HC) was identified, aligned separately and an identity matrix generated.

## Data Availability

The ChIP-seq and microarray data reported in this publication have been deposited in NCBI's Gene Expression Omnibus (Edgar et al., 2002) and are accessible through GEO Series accession numbers GSE154840 and GSE159061, respectively. Additional data that support the findings of this study are available in Supplementary Datasets 1 and 2.

## Acknowledgements

This work received funding from the Swiss National Science Foundation (BSCGI0_157729).

## Author Contributions

E.C. designed and performed experiments, analysed and interpreted data, prepared illustrations and wrote the manuscript. D.K. generated the 3D7/AP2-HC-KO line. R.H.M.C. and C.G.T performed high throughput sequencing, analysed the ChIP-seq data and wrote the corresponding parts of the manuscript and R.B. supervised these experiments, provided resources and wrote the corresponding parts of the manuscript. T.S.V. conceived of the study, designed and supervised experiments, provided resources and wrote the manuscript. All authors contributed to the editing of the manuscript.

