## Supplementary Information for "The ApiAP2 factor PfAP2-HC is an integral component of heterochromatin in the malaria parasite *Plasmodium falciparum*"

This Supplementary Information file includes:

- Supplementary Figures 1-7
- Supplementary Tables 1-2
- Legends pertaining to Supplementary Datasets 1-2

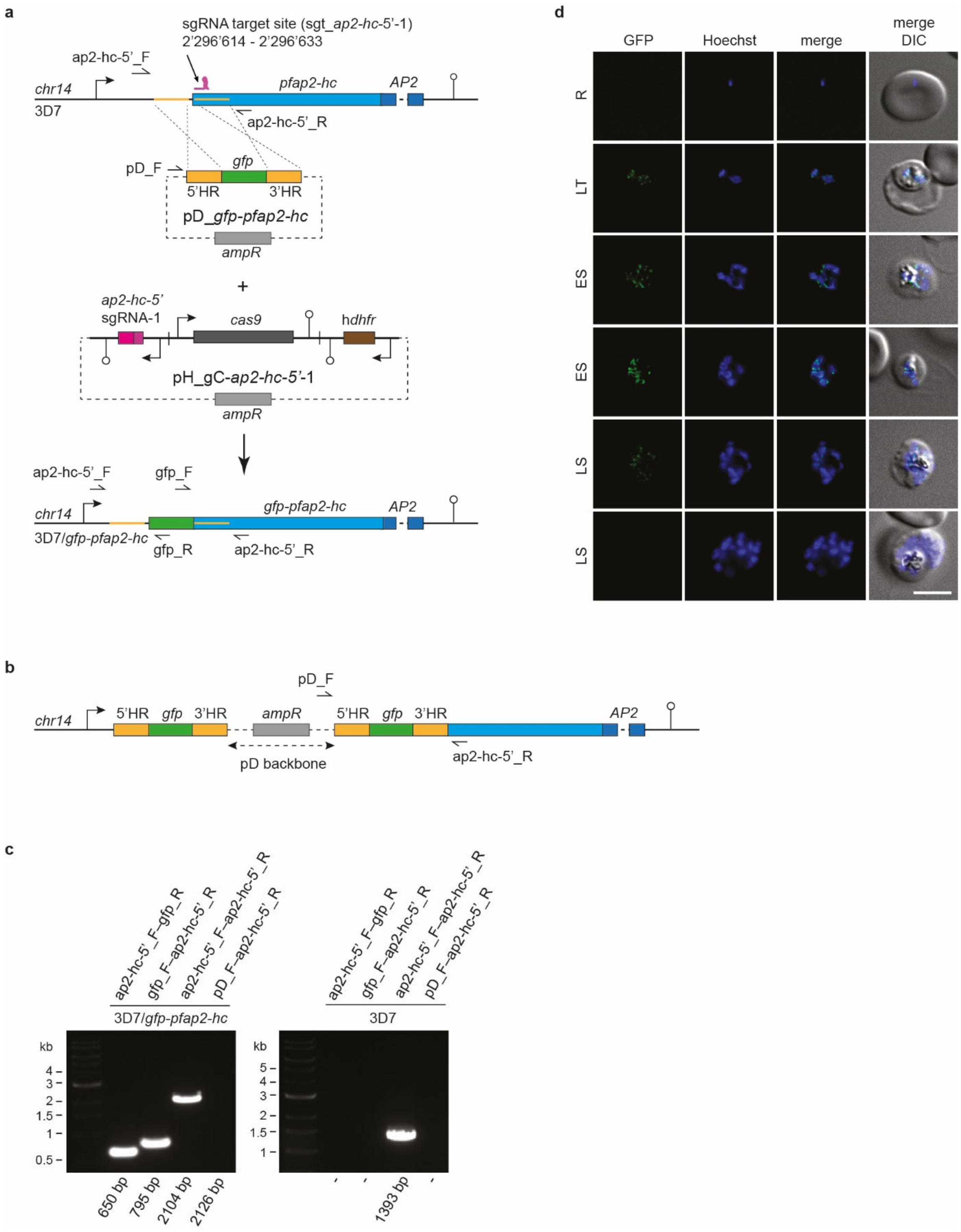


**Supplementary Figure 1. Generation of the 3D7/GFP-PfAP2-HC parasite line and live cell fluorescence imaging. a** Schematic maps of the *pfap2-hc* locus (PF3D7_1456000) in 3D7 parasites (top), the CRISPR/Cas9 transfection vectors pD_*gfp-pfap2-hc* and pH_gC-*ap2-hc-5'*-1 (centre), and the modified *pfap2-hc* locus after CRISPR/Cas9-based genome editing in 3D7/GFP-PfAP2-HC parasites (bottom). The AP2 DBD-encoding sequence, which is interrupted by an intron, is indicated

(AP2, dark blue). The position of the sgt_*ap2-hc-5'*-1 sgRNA target sequence is indicated (chromosome 14 coordinates). The pD_*gfp-pfap2-hc* donor plasmid contains a *gfp* sequence (green) flanked by homology regions (HR, yellow) for homology-directed repair. The pH_gC-*ap2-hc-5'*-1 plasmid contains expression cassettes for SpCas9 (dark grey), the sgRNA (pink) and the h*dhfr* resistance marker (brown). Successful gene editing results in the expression of an N-terminally tagged GFP-PfAP2-HC protein. PCR primer binding sites are indicated by arrows and were used to confirm successful gene editing. **b** Schematic map of the modified *pfap2-hc* locus after CRISPR/Cas9-based genome editing in the event of donor plasmid concatemer integration into the genome. PCR primer binding sites are indicated by arrows and were used to check for donor plasmid concatemer integration. **c** PCR on gDNA from a 3D7/GFP-PfAP2-HC clone and 3D7 wild-type parasites. Primers ap2-hc-5'_F and ap2-hc-5'_R bind to chromosomal sequences outside the HRs and amplify a 2104 bp or 1393 bp fragment from the edited or wild-type *pfap2-hc* locus, respectively. The ap2-hc-5'_F-gfp_R and gfp_F-ap2-hc-5'_R primer combinations are specific for the edited locus and amplify 650 bp and 795 bp fragments, respectively. Primer pD_F binds to the donor plasmid backbone and, when used in combination with primer ap2-hc-5'_R, will amplify a fragment of 2126 bp if a donor plasmid concatemer was integrated into the genome. **d** Live cell fluorescence imaging of 3D7/GFP-PfAP2-HC parasites throughout the IDC. R, ring stage. LT, late trophozoite with two parasites infecting one RBC. ES, early schizont. LS, late schizont. Nuclei were stained with Hoechst. DIC, differential interference contrast. Scale bar, 5 µm.

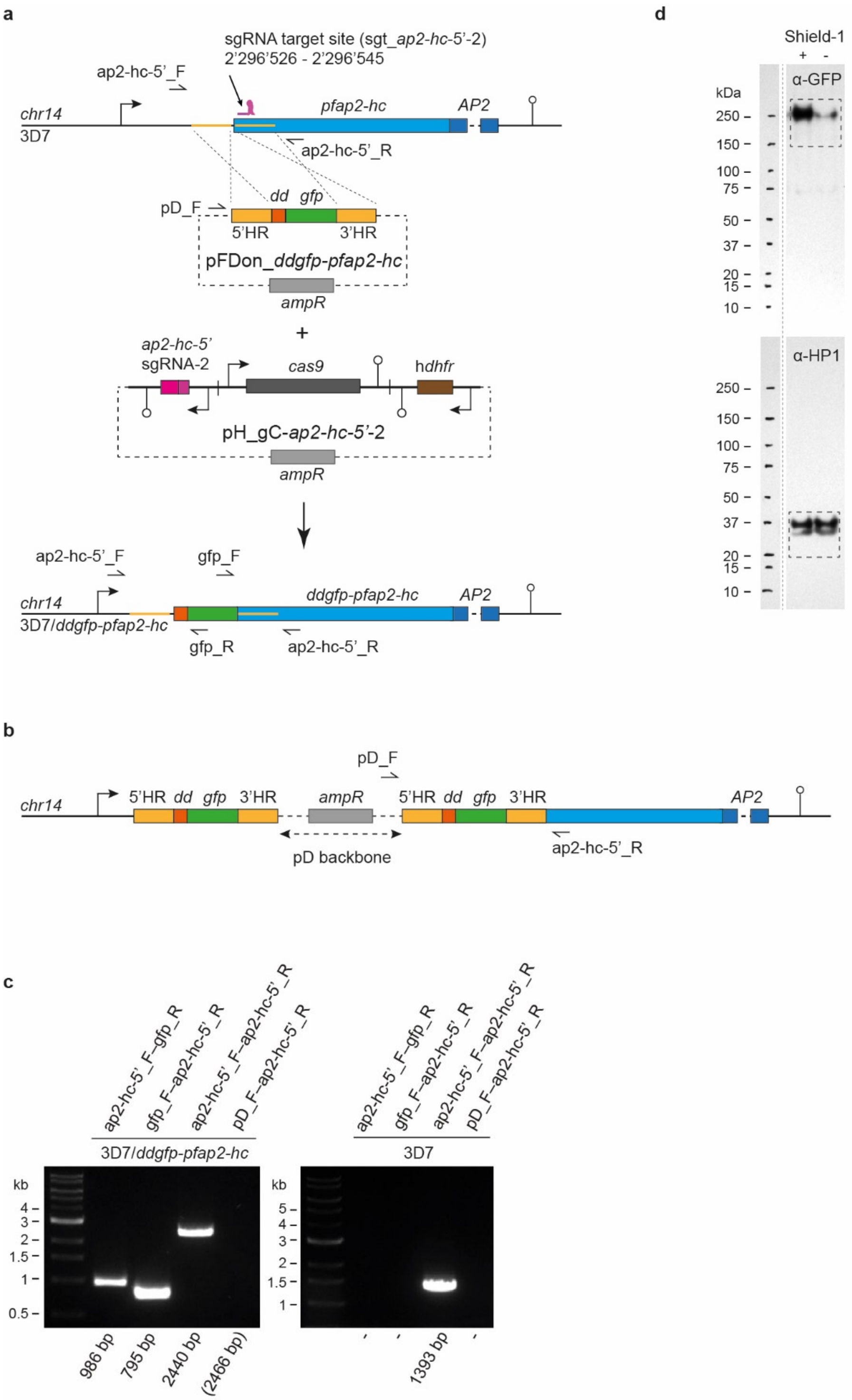


**Supplementary Figure 2. Generation of the 3D7/DDGFP-PfAP2-HC parasite line. a** Schematic maps of the *pfap2-hc* locus (PF3D7_1456000) in 3D7 parasites (top), the CRISPR/Cas9 transfection vectors pFDon_*ddgfp-pfap2-hc* and pH_gC-*ap2-hc-5'*-2 (centre), and the modified *pfap2-hc* locus after CRISPR/Cas9-based genome editing in 3D7/DDGFP-PfAP2-HC parasites (bottom). The AP2

DBD-encoding sequence, which is interrupted by an intron, is indicated (AP2, dark blue). The position of the sgt_*ap2-hc-5'*-1 sgRNA target sequence is indicated (chromosome 14 coordinates). The pFDon_*ddgfp-pfap2-hc* donor plasmid contains an FKBP destabilisation domain (*dd*, orange) and *gfp* sequence (green) flanked by homology regions (HR, yellow) for homology-directed repair. The pH_gC-*ap2-hc-5'*-2 plasmid contains expression cassettes for SpCas9 (dark grey), the sgRNA (pink) and the h*dhfr* resistance marker (brown). Successful gene editing results in the expression of an N-terminally tagged DDGFP-PfAP2-HC protein. PCR primer binding sites are indicated by half arrows and were used to confirm successful gene editing. **b** Schematic map of the modified *pfap2-hc* locus after CRISPR/Cas9-based genome editing in the event of donor plasmid concatemer integration into the genome. PCR primer binding sites are indicated by arrows and were used to check for donor plasmid concatemer integration. **c** PCR on gDNA from a 3D7/DDGFP-PfAP2-HC clone and 3D7 wild-type parasites. Primers ap2-hc-5'_F and ap2-hc-5'_R bind to chromosomal sequences outside the HRs and amplify a 2440 bp or 1393 bp fragment from the edited or wild-type *pfap2-hc* locus, respectively. The ap2-hc-5'_F-gfp_R and gfp_F-ap2-hc-5'_R primer combinations are specific for the edited locus and amplify 986 bp and 795 bp fragments, respectively. Primer pD_F binds to the donor plasmid backbone and, when used in combination with primer ap2-hc-5'_R, will amplify a fragment of 2466 bp if a donor plasmid concatemer was integrated into the genome. **d** Full sized Western blot of the sections shown in Fig. 2b showing DDGFP-PfAP2-HC expression levels in 3D7/DDGFP-PfAP2-HC parasites grown in the presence (+) or absence (-) of Shield-1. The membrane was first probed with α-GFP antibodies (top) before inactivation of horseradish peroxidase with 2 mM $NaN_3$, followed by re-probing with the α-PfHP1 antibodies (bottom) used as a loading control. Dashed boxes show the sections presented in Fig. 2b.

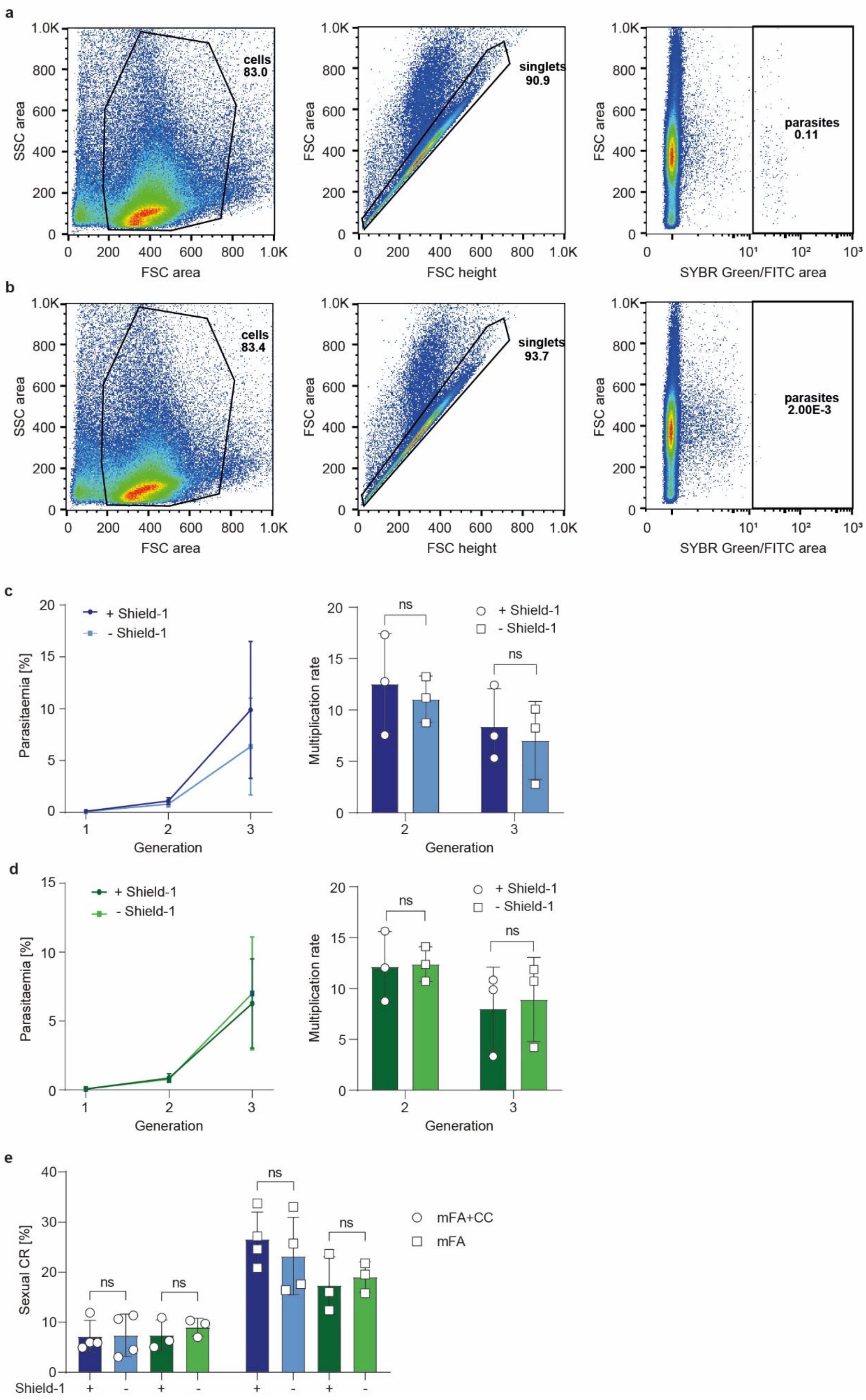
a
cells 83.0
singlets 90.9
parasites 0.11
SSC area
FSC area
FSC height
SYBR Green/FITC area
b
cells 83.4
singlets 93.7
parasites 2.00E-3
c
+ Shield-1
- Shield-1
Parasitaemia [%]
Multiplication rate
Generation
ns
d
e
Sexual CR [%]
mFA+CC
mFA
Shield-1

**Supplementary Figure 3. Multiplication rates and gametocyte conversion rates of 3D7/DDGFP-PfAP2-HC parasites. a**, **b** Gating strategy applied to flow cytometry data obtained from multiplication assays. Representative flow cytometry plots of an infected (3D7/WT, - Shield-1, panel a) and uninfected RBC control sample (panel b) measured on day 1 of the multiplication assay. The first plot shows the gate to remove debris smaller than cell size to include only the 'cells' population. The second plot shows the gate to include only single measurement events, termed 'singlets', and the third gate separates uninfected from infected RBCs based on the SYBR Green intensity of the uninfected RBC control, termed 'parasites'. The numbers are the percentage of events included within the gate, with the final gate 'parasites' reflecting the parasitaemia of the sample. This gating strategy was applied to all flow cytometry data shown in panels c and d, and in Supplementary Fig. 4. **c**, **d** Flow cytometry data showing the increase in parasitaemia (left) and parasite multiplication rates (right) in two subsequent generations of 3D7/DDGFP-PfAP2-HC (panel c) parasites grown in the presence (+, dark blue) or absence (-, light blue) of Shield-1 and 3D7/WT (panel d) parasites grown in the presence (+, dark green) or absence (-, light green) of Shield-1. The mean ±SD of three biological replicates are shown. Data points of individual replicates are shown for parasite multiplication rates and represented by open circles (+ Shield-1) or open squares (- Shield-1). ns, not significant (paired two-tailed Student's t test). **e** Sexual conversion rates of 3D7/DDGFP-PfAP2-HC (dark/light blue) and 3D7/WT (dark/light green) parasites exposed to mFA+CC medium (conditions inhibiting sexual commitment, open circles) or mFA medium (conditions inducing sexual commitment, open squares)[1]. Parasites grown in the presence (+) or absence (-) of Shield-1 are compared. The mean ±SD of four biological replicates of 3D7/DDGFP-PfAP2-HC and three biological replicates of 3D7/WT are shown. Data points of individual replicates are shown. ns, not significant (paired two-tailed Student's t test). CR, conversion rate.

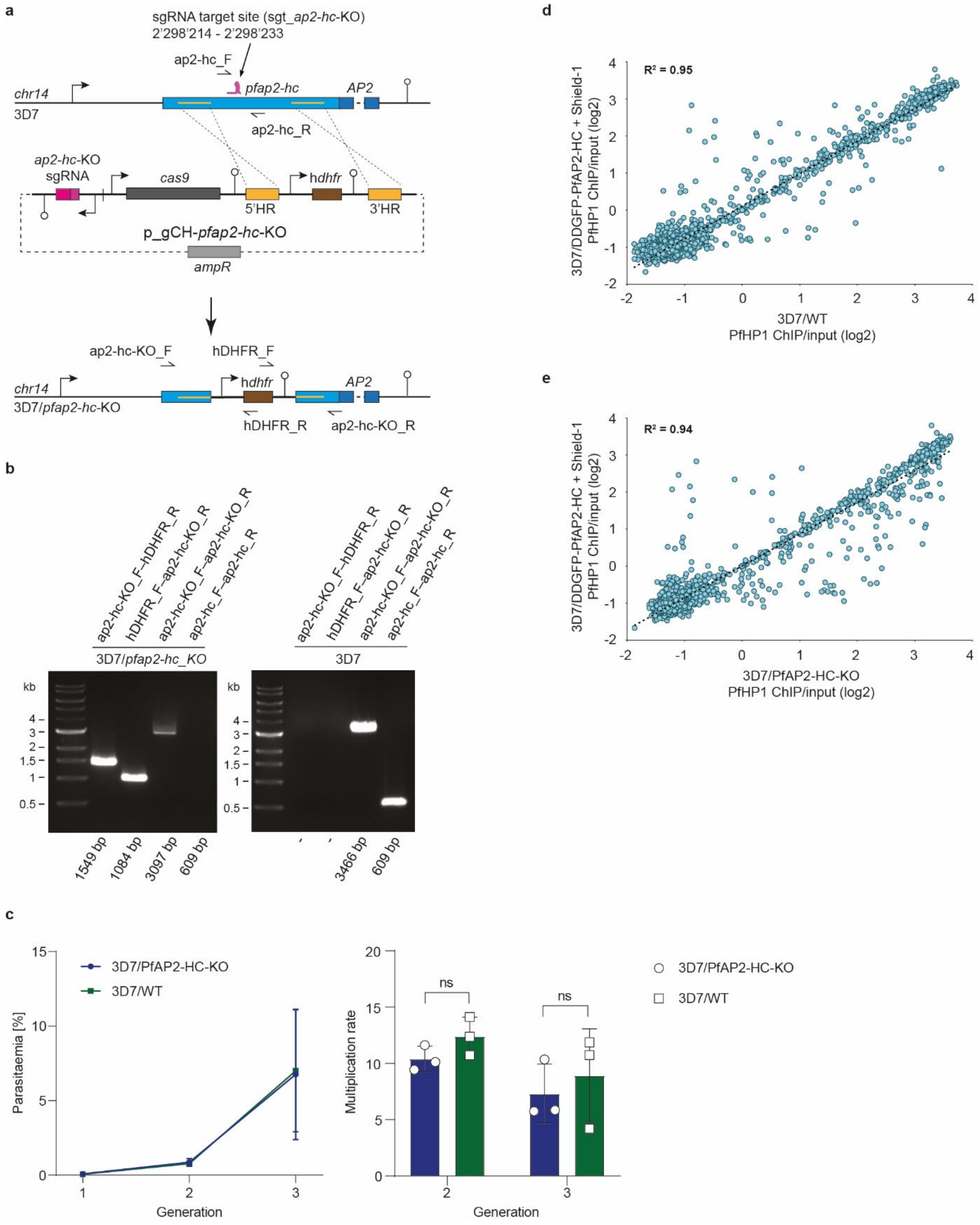


**Supplementary Figure 4. Generation of the 3D7/PfAP2-HC-KO parasite line. a** Schematic maps of the *pfap2-hc* locus (PF3D7_1456000) in 3D7 parasites (top), the p_gCH-*pfap2-hc*-KO transfection vector (centre), and the modified *pfap2-hc* locus after CRISPR/Cas9-based genome editing in 3D7/PfAP2-HC-KO parasites (bottom). The AP2 DBD-encoding sequence, which is interrupted by an intron, is indicated (AP2, dark blue). The position of the sgt_*ap2-hc*-KO sgRNA target sequence is indicated (chromosome 14 coordinates). The p_gCH-*pfap2-hc*-KO plasmid contains expression cassettes for SpCas9 (dark grey), the sgRNA (pink) and the h*dhfr* resistance marker (brown) flanked by two homology regions (HR, yellow) for homology-directed repair. Successful gene editing results in the h*dhfr* expression cassette replacing a section of the *pfap2-hc* gene, disrupting its expression. PCR primer binding sites are indicated by arrows and were used to confirm successful gene editing. **b** PCR on gDNA from 3D7/PfAP2-HC-KO and 3D7 wild-type parasites. Primers ap2-hc-KO_F and ap2-hc-KO_R bind to chromosomal sequences outside the HRs and amplify a 3097 bp or 3466 bp fragment from the edited or wild-type *pfap2-hc* locus, respectively. The ap2-hc-KO_F-hDHFR_R and hDHFR_F-ap2-hc-KO_R primer combinations are specific for the edited locus and amplify 1549 bp and 1084 bp fragments, respectively. Primer combination ap2-hc_F-ap2-hc_R is specific for the wild-type locus and amplifies a 609 bp fragment. **c** Flow cytometry data showing the increase in parasitaemia (left) and parasite multiplication rates (right) in two subsequent generations of 3D7/PfAP2-HC-KO (dark blue) and 3D7/WT (dark green) parasites. The 3D7/WT data is identical to those shown in Supplementary Fig. 2d (3D7/WT grown in the absence of Shield-1). The mean ±SD of three biological replicates are shown. Data points of individual replicates are shown for parasite multiplication rates and represented by open circles (3D7/PfAP2-HC-KO) or open squares (3D7/WT). ns, not significant (paired two-tailed Student's t test). **d, e** Scatterplots of average log2-transformed α-PfHP1 ChIP/input values for all parasite genes in 3D7/WT and 3D7/DDGFP-PfAP2-HC schizonts grown in the presence (+) of Shield-1 (panel c) and in 3D7/PfAP2-HC-KO and 3D7/DDGFP-PfAP2-HC schizonts grown in the presence (+) of Shield-1 (panel d). Depicted regression lines are based on heterochromatic genes only (log2 ratio α-PfHP1/input ≥ 0). The coefficients of determination ($R^2$) are shown on the top left.

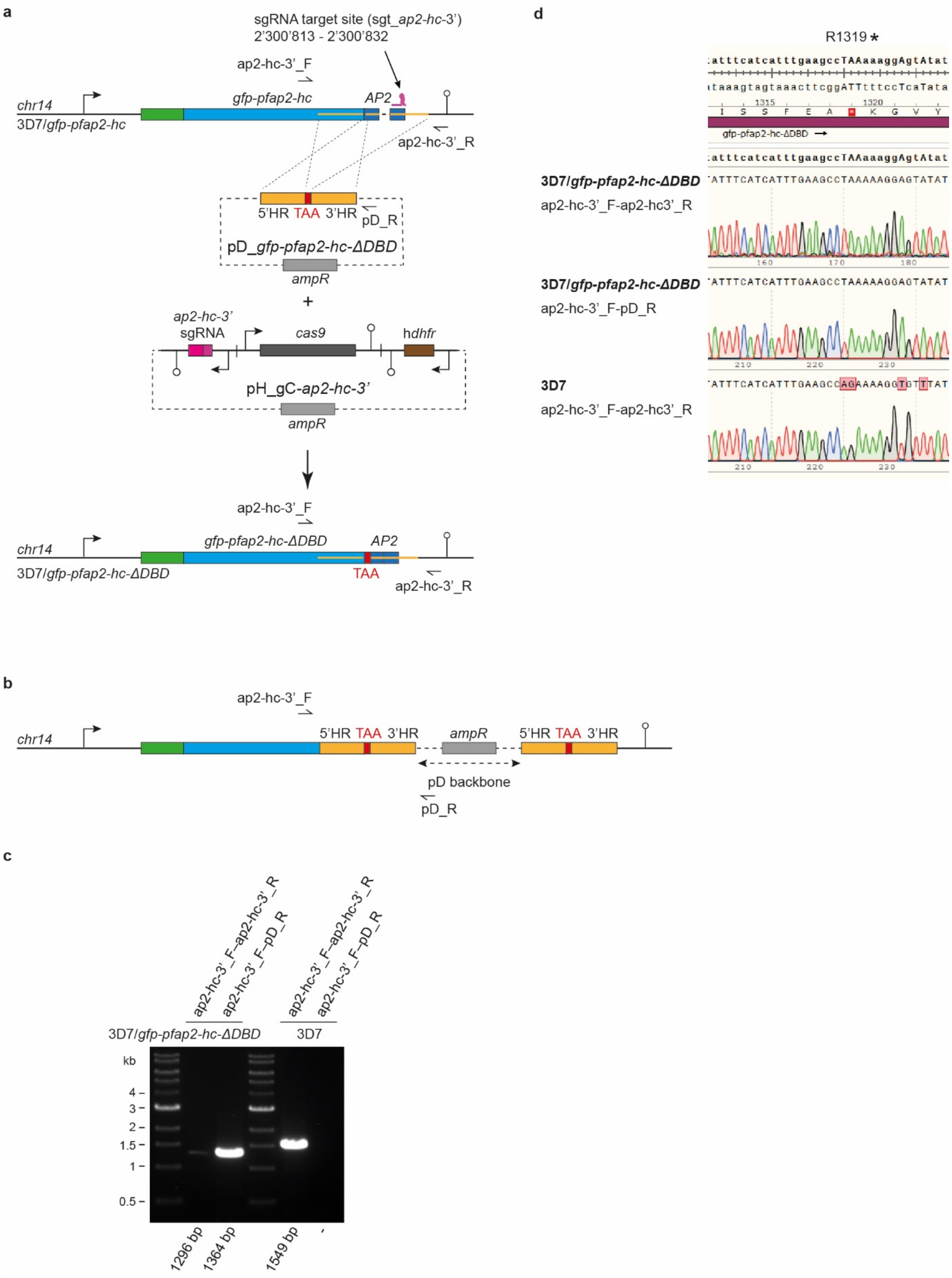


**Supplementary Figure 5. Generation of the 3D7/GFP-PfAP2-HC-ΔDBD parasite line. a**

Schematic maps of the *gfp-pfap2-hc* locus in 3D7/GFP-PfAP2-HC parasites (top, see Supplementary Fig. 1), the CRISPR/Cas9 transfection vectors pD_*gfp-pfap2-hc-ΔDBD* and pH_gC-*ap2-hc-3'* (centre), and the modified *gfp-pfap2-hc* locus after CRISPR/Cas9-based genome editing in 3D7/GFP-PfAP2-HC-ΔDBD parasites (bottom). The AP2 DBD-encoding sequence, which is interrupted by an intron, is indicated (AP2, dark blue). The position of the sgt_*ap2-hc-3'* sgRNA target sequence is indicated (chromosome 14 coordinates). The pD_*gfp-pfap2-hc-ΔDBD* donor plasmid contains a premature TAA stop codon (red) flanked by homology regions (HR, yellow) for homology-directed repair. The pH_gC-*ap2-hc-3'* plasmid contains expression cassettes for SpCas9 (dark grey), the sgRNA (pink) and the h*dhfr* resistance marker (brown). Successful gene editing results in the expression of a truncated GFP-PfAP2-HC protein lacking the AP2 DNA-binding domain (GFP-PfAP2-HC-ΔDBD). PCR primer binding sites are indicated by arrows and were used to confirm successful gene editing. **b** Schematic map of the modified *gfp-pfap2-hc* locus after CRISPR/Cas9-based genome editing in the event of donor plasmid concatemer integration into the genome. PCR primer binding sites are indicated by arrows and were used to check for donor plasmid concatemer integration. **c** PCR on gDNA from 3D7/GFP-PfAP2-HC-ΔDBD and 3D7 wild-type parasites. Primers ap2-hc-3'_F and ap2-hc-3'_R bind to chromosomal sequences outside the HRs and amplify a 1296 bp or 1549 bp fragment from the edited or wild-type *pfap2-hc* locus, respectively. Primer pD_R binds to the donor plasmid backbone and, when used in combination with primer ap2-hc-3'_F, amplifies a fragment of 1364 bp if a donor plasmid concatemer was integrated into the genome. **d** Sanger sequencing of the two PCR products ap2-hc-3'_F-ap2-hc-3'_R (top) and ap2-hc-3'_F-pD_R (middle) amplified from 3D7/GFP-PfAP2-HC-ΔDBD parasites (see panel b, lanes 2 and 3) confirms the successful introduction of the AG→TA double mutation creating a premature STOP codon (R1319*). The PCR product ap2-hc-3'_F-ap2-hc-3'_R (bottom) amplified from 3D7 wild-type parasites (see panel b, lane 5) shows the wild-type sequence. Additional mutations downstream of the AG→TA double mutation are part of the re-codonised sequence introduced to avoid homologues recombination at an undesired location to ensure correct CRISPR/Cas9 genome editing.

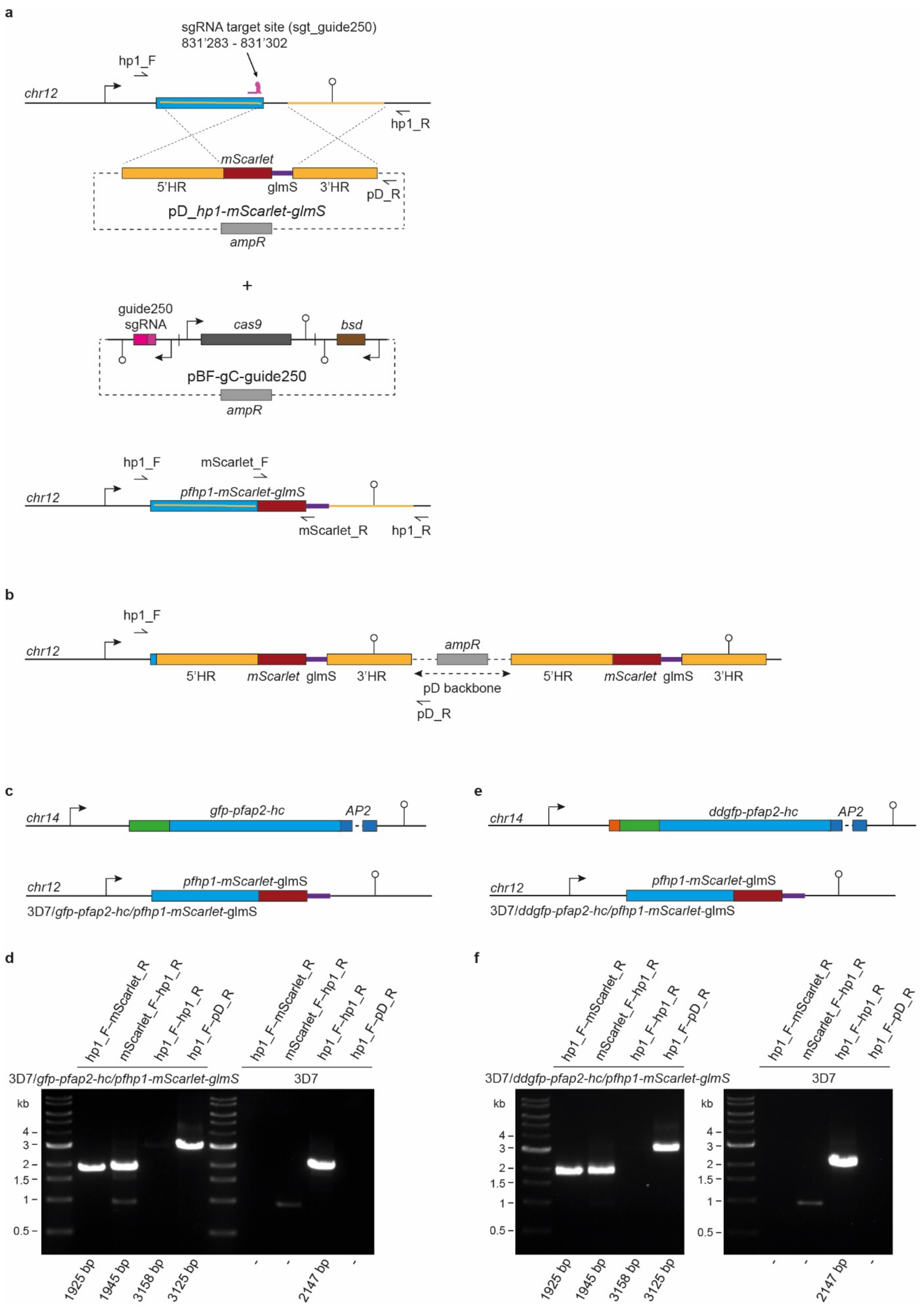


**Supplementary Figure 6. Generation of the 3D7/GFP-PfAP2-HC/PfHP1-mScarlet-glmS and 3D7/DDGFP-PfAP2-HC/PfHP1-mScarlet-glmS parasite lines. a** Schematic maps of the wild-type *pfhp1* locus (PF3D7_1220900) in 3D7/GFP-PfAP2-HC and 3D7/DDGFP-PfAP2-HC parasites (top), the CRISPR/Cas9 transfection vectors pD_*hp1-mScarlet-glmS* and pBF-gC-guide250[2] (centre), and the modified *pfhp1* locus after CRISPR/Cas9-based genome editing (bottom). The position of the sgt_guide250 sgRNA target sequence is indicated (chromosome 12 coordinates). The pD_*hp1-mScarlet-glmS* donor plasmid contains the *mScarlet* sequence (*red*) followed by the *glmS* ribozyme sequence (purple) flanked by homology regions (HR, yellow) for homology-directed repair. The pBF-gC-guide250 plasmid[2] contains expression cassettes for SpCas9 (dark grey), the sgRNA (pink) and the blasticidin deaminase (*bsd*) resistance marker (brown). Successful gene editing results in the expression of a C-terminally tagged PfHP1-mScarlet protein controlled by the *glmS* ribozyme element. PCR primer binding sites are indicated by half arrows and were used to confirm successful gene editing. **b** Schematic map of the modified *pfhp1* locus after CRISPR/Cas9-based genome editing in the event of donor plasmid concatemer integration into the genome. PCR primer binding sites are indicated by arrows and were used to check for donor plasmid concatemer integration. **c** Schematic maps of the *gfp-pfap2-hc* locus (top, see Supplementary Fig. 1) and the *pfhp1-mScarlet-glmS* locus (bottom) in successfully edited 3D7/GFP-PfAP2-HC/PfHP1-mScarlet-glmS parasites. **d** PCR on gDNA from 3D7/GFP-PfAP2-HC/PfHP1-mScarlet-glmS and 3D7 wild-type parasites. Primers hp1_F and hp1_R bind to chromosomal sequences outside the HRs and amplify a 3158 bp or 2147 bp fragment from the edited or wild-type *pfhp1* locus, respectively. The hp1_F-mScarlet_R and mScarlet_F-hp1_R primer combinations are specific for the edited locus and amplify 1925 bp and 1945 bp fragments, respectively. Primer pD_R binds to the donor plasmid backbone and, when used in combination with primer hp1_F, will amplify a fragment of 3125 bp if a donor plasmid concatemer was integrated into the genome. **e** Schematic maps of the *ddgfp-pfap2-hc* locus (top, see Supplementary Fig. 2) and the *pfhp1-mScarlet-glmS* locus (bottom) in successfully edited 3D7/DDGFP-PfAP2-HC/PfHP1-mScarlet-glmS parasites. **f** PCR on gDNA from 3D7/DDGFP-PfAP2-HC/PfHP1-mScarlet-glmS and 3D7 wild-type parasites. Primer explanations are as in panel d. Primer combination mScarlet_F-hp1_R results in a faint non-specific product at ~1000 bp in all reactions (panels d and f).

a

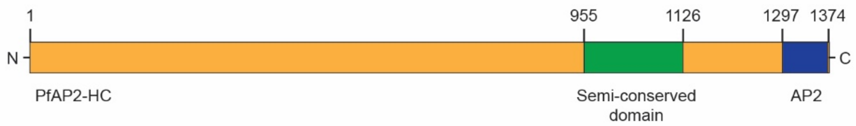


b

| 1 – 1374: Full length, % sequence identity | | | | | | | | |
|---|---|---|---|---|---|---|---|---|
| | **Pv** | **Pk** | **Poc** | **Pm** | **Pb** | **Py** | **Pc** | **Pf** |
| **Pv** | 100 | 81 | 46 | 46 | 37 | 38 | 37 | 35 |
| **Pk** | 81 | 100 | 47 | 46 | 37 | 37 | 36 | 35 |
| **Poc** | 46 | 47 | 100 | 46 | 37 | 38 | 38 | 31 |
| **Pm** | 46 | 46 | 46 | 100 | 40 | 41 | 40 | 34 |
| **Pb** | 37 | 37 | 37 | 40 | 100 | 91 | 83 | 32 |
| **Py** | 38 | 37 | 38 | 41 | 91 | 100 | 85 | 33 |
| **Pc** | 37 | 36 | 38 | 40 | 83 | 85 | 100 | 31 |
| **Pf** | 35 | 35 | 31 | 34 | 32 | 33 | 31 | 100 |

c

| 955 – 1126: Semi-conserved domain, % sequence identity | | | | | | | | |
|---|---|---|---|---|---|---|---|---|
| | **Pm** | **Poc** | **Pv** | **Pk** | **Pf** | **Pb** | **Py** | **Pc** |
| **Pm** | 100 | 78 | 77 | 76 | 64 | 62 | 64 | 62 |
| **Poc** | 78 | 100 | 76 | 75 | 67 | 59 | 60 | 60 |
| **Pv** | 77 | 76 | 100 | 95 | 66 | 58 | 60 | 59 |
| **Pk** | 76 | 75 | 95 | 100 | 66 | 59 | 60 | 59 |
| **Pf** | 64 | 67 | 66 | 66 | 100 | 53 | 56 | 55 |
| **Pb** | 62 | 59 | 58 | 59 | 53 | 100 | 95 | 92 |
| **Py** | 64 | 60 | 60 | 60 | 56 | 95 | 100 | 95 |
| **Pc** | 62 | 60 | 59 | 59 | 55 | 92 | 95 | 100 |

d

| 1297 – 1374: AP2 domain, % sequence identity | | | | | | | | |
|---|---|---|---|---|---|---|---|---|
| | **Pb** | **Pc** | **Py** | **Pm** | **Poc** | **Pv** | **Pk** | **Pf** |
| **Pb** | 100 | 99 | 99 | 94 | 94 | 95 | 95 | 90 |
| **Pc** | 99 | 100 | 100 | 95 | 95 | 96 | 96 | 91 |
| **Py** | 99 | 100 | 100 | 95 | 95 | 96 | 96 | 91 |
| **Pm** | 94 | 95 | 95 | 100 | 97 | 99 | 99 | 94 |
| **Poc** | 94 | 95 | 95 | 97 | 100 | 96 | 96 | 94 |
| **Pv** | 95 | 96 | 96 | 99 | 96 | 100 | 100 | 95 |
| **Pk** | 95 | 96 | 96 | 99 | 96 | 100 | 100 | 95 |
| **Pf** | 90 | 91 | 91 | 94 | 94 | 95 | 95 | 100 |

**Supplementary Figure 7. AP2-HC amino acid sequence comparison between orthologs of different *Plasmodium* species. a** Schematic map of the PfAP2-HC protein showing the location of the AP2 domain (blue) and a semi-conserved domain (green) identified via a multiple sequence alignment of AP2-HC orthologues from *P. vivax* (PVX_117665), *P. knowlesi* (PKNH_1225800), *P. malariae* (PmUG01_12060900), *P. ovale curtisi* (PocGH01_12058800), *P. berghei* (PBANKA_1319700), *P. yoelii* (PY17X_1323500) and *P. chabaudi* (PCHAS_1323000). Numbers refer to the amino acid position within the PfAP2-HC sequence. **b**, **c**, **d** Amino acid sequence identity matrices of AP2-HC orthologues from eight *Plasmodium* species, comparing the full length protein (panel b), a semi-conserved domain of 172 amino acids (panel c), and the AP2 domain (panel d). Pf, *P. falciparum*. Pv, *P. vivax*. Pk, *P. knowlesi*. Pm, *P. malariae*. Poc, *P. ovale curtisi*. Pb, *P. berghei*. Py, *P. yoelii*. Pc, *P. chabaudi*.

**Supplementary Table 1. Oligonucleotide sequences used for cloning of CRISPR/Cas9 transfection vectors.** Oligonucleotide names and sequences are shown alongside the plasmid and parasite cell lines they were used to generate. Oligonucleotide sequences used to generate PCR fragments for Gibson assembly reactions (Gibson overhangs) are in upper case. Oligonucleotide sequences required for ligation of annealed double-stranded sgRNA-encoding sequences into the *Bsa*I site of the sgRNA expression cassette are italicized in upper case. A premature STOP codon is highlighted in red font.

| **Oligonucleotide name** | **Oligonucleotide sequence 5'→3'** | **Plasmid name** | **Cell line name** |
|---|---|---|---|
| PCRA_F[3] | ctggcgtaatagcgaagagg | pD_*gfp-pfap2-hc,*<br>pD_*gfp-pfap2-hc-ΔDBD* | 3D7/GFP-PfAP2-HC,<br>3D7/GFP-PfAP2-HC-ΔDBD |
| PCRA_R[3] | cattaatgaatcggccaacg | pD_*gfp-pfap2-hc,*<br>pD_*gfp-pfap2-hc-ΔDBD* | 3D7/GFP-PfAP2-HC,<br>3D7/GFP-PfAP2-HC-ΔDBD |
| ap2-hc-5'_HR1_F | CGTTGGCCGATTCATTAATGcttatatttgtattcagttgattctaac | pD_*gfp-pfap2-hc* | 3D7/GFP-PfAP2-HC |
| ap2-hc-5'_HR1_R | TTCTCCTTTACTCATattttattcttattttgtgtattggttataag | pD_*gfp-pfap2-hc* | 3D7/GFP-PfAP2-HC |
| ap2-hc-5'_GFP_F | AATAAGAATAAAATatgagtaaaggagaagaacttttcac | pD_*gfp-pfap2-hc* | 3D7/GFP-PfAP2-HC |
| ap2-hc-5'_GFP_R | ACTGAATATTCATTtttgtatagttcatccatgccatg | pD_*gfp-pfap2-hc* | 3D7/GFP-PfAP2-HC |
| ap2-hc-5'_HR2_re_F | TGAACTATACAAAaatgaatattcagtttataataagcc | pD_*gfp-pfap2-hc* | 3D7/GFP-PfAP2-HC |
| ap2-hc-5'_HR2_re_R | CCTCTTCGCTATTACGCCAGggtcatctaaatttccattagg | pD_*gfp-pfap2-hc* | 3D7/GFP-PfAP2-HC |
| sgRNA_ap2-hc-5'-1_F | *TATT*gaaacacataacgagcttaa | pH_gC-*ap2-hc-5'*-1 | 3D7/GFP-PfAP2-HC |
| sgRNA_ap2-hc-5'-1_R | *AAAC*ttaagctcgttatgtgtttc | pH_gC-*ap2-hc-5'*-1 | 3D7/GFP-PfAP2-HC |
| ap2-hc-5'_HR3_F | CGAGTCAGTGAGCGAGGActtatatttgtattcagttgattctaac | pFDon_*ddgfp-pfap2-hc* | 3D7/DDGFP-PfAP2-HC |
| ap2-hc-5'_HR3_R | GTTTCCACCTGCACTCCCATattttattcttattttgtgtattggttataag | pFDon_*ddgfp-pfap2-hc* | 3D7/DDGFP-PfAP2-HC |
| ap2-hc-5'_DD_F | ACACAAAATAAGAATAAAATatgggagtgcaggtggaaac | pFDon_*ddgfp-pfap2-hc* | 3D7/DDGFP-PfAP2-HC |
| ap2-hc-5'_DD_R | TTCTTCTCCTTTACTCATACTAGAACCGGTttccagttttagaagctccacac | pFDon_*ddgfp-pfap2-hc* | 3D7/DDGFP-PfAP2-HC |
| ap2-hc-5'_HR4_F | GAGCTTCTAAAACTGGAAACCGGTTCTAGTatgagtaaaggagaagaacttttcac | pFDon_*ddgfp-pfap2-hc* | 3D7/DDGFP-PfAP2-HC |
| ap2-hc-5'_HR4_R | CTTTTCTCTTGTGGATCCGggtcatctaaatttccattagg | pFDon_*ddgfp-pfap2-hc* | 3D7/DDGFP-PfAP2-HC |
| sgRNA_ap2-hc-5'-2_F | *TATT*taacaatatttctgtatcta | pH_gC-*ap2-hc-5'*-2 | 3D7/DDGFP-PfAP2-HC |
| sgRNA_ap2-hc-5'-2_R | *AAAC*tagatacagaaatattgtta | pH_gC-*ap2-hc-5'*-2 | 3D7/DDGFP-PfAP2-HC |
| ap2-hc-3'_HR1_re_F | CGTTGGCCGATTCATTAATGtaatagagatgaaaatagacaggca | pD_*gfp-pfap2-hc-ΔDBD* | 3D7/GFP-PfAP2-HC-ΔDBD |
| ap2-hc-3'_HR1_re_R | acttttatcataatatactcctttTTAggcttcaaatgatgaaataaccc | pD_*gfp-pfap2-hc-ΔDBD* | 3D7/GFP-PfAP2-HC-ΔDBD |
| ap2-hc-3'_HR2_re_F | ggttatttcatcatttgaagccTAAaaaggagtatattatgataaaagtag | pD_*gfp-pfap2-hc-ΔDBD* | 3D7/GFP-PfAP2-HC-ΔDBD |
| ap2-hc-3'_HR2_re_R | aacaataattcagcttctttttctctctccattctattgctttttgtc | pD_*gfp-pfap2-hc-ΔDBD* | 3D7/GFP-PfAP2-HC-ΔDBD |
| ap2-hc-3'_HR3_F | acaaaaagcaatagaatggagagagaaaaaagaagctgaattattgtt | pD_*gfp-pfap2-hc-ΔDBD* | 3D7/GFP-PfAP2-HC-ΔDBD |
| ap2-hc-3'_HR3_R | CCTCTTCGCTATTACGCCAGatcatatccatctccatacacaaatg | pD_*gfp-pfap2-hc-ΔDBD* | 3D7/GFP-PfAP2-HC-ΔDBD |

| | | | |
|---|---|---|---|
| sgRNA_ap2-hc-3'_F | *TATT*ctagacaaaaggctattgaa | pH_gC-*ap2-hc-3'* | 3D7/GFP-PfAP2-HC-ΔDBD |
| sgRNA_ap2-hc-3'_R | *AAAC*ttcaatagccttttgtctag | pH_gC-*ap2-hc-3'* | 3D7/GFP-PfAP2-HC-ΔDBD |
| ap2-hc-KO_HR1_F | TCAGGGTAGCTGATATCGGA TCCcacataacgagcttaatgg | p_gCH-*pfap2-hc*-KO | 3D7/PfAP2-HC-KO |
| ap2-hc-KO_HR1_R | CCTTTTCTCTTGTcattatattcttca atgtctattac | p_gCH-*pfap2-hc*-KO | 3D7/PfAP2-HC-KO |
| ap2-hc-KO_hDHFR_F | AAGAATATAATGacaagagaaaa ggcagaaac | p_gCH-*pfap2-hc*-KO | 3D7/PfAP2-HC-KO |
| ap2-hc-KO_hDHFR_R | CATTACACAAGGACtttaataaata tgttcttatatataatgag | p_gCH-*pfap2-hc*-KO | 3D7/PfAP2-HC-KO |
| ap2-hc-KO_HR2_F | CATATTTATTAAAgtccttgtgtaat gaaaatatc | p_gCH-*pfap2-hc*-KO | 3D7/PfAP2-HC-KO |
| ap2-hc-KO_HR2_R | GAGCGAGGAAGCGGAAGCT Tgtgtacttggttgatcatatag | p_gCH-*pfap2-hc*-KO | 3D7/PfAP2-HC-KO |
| sgRNA_ap2-hc-KO_F | *TATT*cgttgtactagtaacattgg | p_gCH-*pfap2-hc*-KO | 3D7/PfAP2-HC-KO |
| sgRNA_ap2-hc-KO_R | *AAAC*ccaatgttactagtacaacg | p_gCH-*pfap2-hc*-KO | 3D7/PfAP2-HC-KO |
| F158[2] | CGTTGGCCGATTCATTAATG aaaggatattcagatgatgag | pD_*hp1-mScarlet-glmS* | 3D7/GFP-PfAP2-HC/PfHP1-mScarlet-glmS, 3D7/DDGFP-PfAP2-HC/PfHP1-mScarlet-glmS |
| hp1_HR1_R | CCTTTACTACCTGCGGATCC cgctgttctatatcttaatc | pD_*hp1-mScarlet-glmS* | 3D7/GFP-PfAP2-HC/PfHP1-mScarlet-glmS, 3D7/DDGFP-PfAP2-HC/PfHP1-mScarlet-glmS |
| hp1_mScarlet_F | GATTAAGATATAGAACAGC Gggatccgcaggtagtaaagg | pD_*hp1-mScarlet-glmS* | 3D7/GFP-PfAP2-HC/PfHP1-mScarlet-glmS, 3D7/DDGFP-PfAP2-HC/PfHP1-mScarlet-glmS |
| hp1_mScarlet_R | TTGAGAAAATAAGAACAAG Atcatttatataattcatccattcc | pD_*hp1-mScarlet-glmS* | 3D7/GFP-PfAP2-HC/PfHP1-mScarlet-glmS, 3D7/DDGFP-PfAP2-HC/PfHP1-mScarlet-glmS |
| hp1_glmS_F | GAATGGATGAATTATATAA ATGAtcttgttcttattttctcaatagg | pD_*hp1-mScarlet-glmS* | 3D7/GFP-PfAP2-HC/PfHP1-mScarlet-glmS, 3D7/DDGFP-PfAP2-HC/PfHP1-mScarlet-glmS |
| hp1_glmS_R | TGTATATTTGCATAATAAAA atttttcttcctcctaagattgtaaaag | pD_*hp1-mScarlet-glmS* | 3D7/GFP-PfAP2-HC/PfHP1-mScarlet-glmS, 3D7/DDGFP-PfAP2-HC/PfHP1-mScarlet-glmS |
| hp1_HR2_F | ATCTTAGGAGGAAGAAAAA Ttttttattatgcaaatatacatatatac | pD_*hp1-mScarlet-glmS* | 3D7/GFP-PfAP2-HC/PfHP1-mScarlet-glmS, 3D7/DDGFP-PfAP2-HC/PfHP1-mScarlet-glmS |
| R163[2] | CCTCTTCGCTATTACGCCAG gaggttaaaattctaaactatatg | pD_*hp1-mScarlet-glmS* | 3D7/GFP-PfAP2-HC/PfHP1-mScarlet-glmS, 3D7/DDGFP-PfAP2-HC/PfHP1-mScarlet-glmS |

**Supplementary Table 2. Primers used for PCRs on gDNA of CRISPR/Cas9-edited gene loci.**

Primer names and sequences are shown alongside the parasite cell lines from which gDNA was extracted to carry out PCRs to confirm successful gene editing.

| Primer name | Primer sequence 5' → 3' | Cell line name |
|---|---|---|
| pD_F | accgcctttgagtgagc | |
| pD_R | cgaaaagtgccacctgacg | |
| ap2-hc-5'_F | attacttatattttatttttctcttcaaagaaa | 3D7/GFP-PfAP2-HC, 3D7/DDGFP-PfAP2-HC |
| gfp_R | tccagtgaaaagttcttctcct | 3D7/GFP-PfAP2-HC, 3D7/DDGFP-PfAP2-HC |
| gfp_F | acatggcatggatgaactatacaaa | 3D7/GFP-PfAP2-HC, 3D7/DDGFP-PfAP2-HC |
| ap2-hc-5'_R | acacaaacgctttcactatcttct | 3D7/GFP-PfAP2-HC, 3D7/DDGFP-PfAP2-HC |
| ap2-hc-KO_F | ctaacaatatttctgtatctaaggg | 3D7/PfAP2-HC-KO |
| hDHFR_R | aacgatgcagtttagcgaacc | 3D7/PfAP2-HC-KO |
| hDHFR_F | atgtccaggaggagaaagg | 3D7/PfAP2-HC-KO |
| ap2-hc-KO_R | aggttattttttaactgattatttagagg | 3D7/PfAP2-HC-KO |
| ap2-hc_F | attaaagaattttggagttcctcc | 3D7/PfAP2-HC-KO |
| ap2-hc_R | ctttgtgcattcatcctcagg | 3D7/PfAP2-HC-KO |
| ap2-hc-3'_F | aataaccttcagaagaaatcgcaaa | 3D7/GFP-PfAP2-HC-ΔDBD |
| ap2-hc-3'_R | atcggatatttattctcttgctgttg | 3D7/GFP-PfAP2-HC-ΔDBD |
| hp1_F | gtgtgtgtttaagaaaaaaatatg | 3D7/GFP-PfAP2-HC/PfHP1-mScarlet-glmS, 3D7/DDGFP-PfAP2-HC/PfHP1-mScarlet-glmS |
| mScarlet_R | tgtatatttgcataataaaatcatttatataattcatccattccacc | 3D7/GFP-PfAP2-HC/PfHP1-mScarlet-glmS, 3D7/DDGFP-PfAP2-HC/PfHP1-mScarlet-glmS |
| mScarlet_F | gattaagatatagaacagcgggatccgcaggtagtaaagg | 3D7/GFP-PfAP2-HC/PfHP1-mScarlet-glmS, 3D7/DDGFP-PfAP2-HC/PfHP1-mScarlet-glmS |
| hp1_R | catgtagccaaaatatgtg | 3D7/GFP-PfAP2-HC/PfHP1-mScarlet-glmS, 3D7/DDGFP-PfAP2-HC/PfHP1-mScarlet-glmS |

**Supplementary Dataset 1. ChIP-seq enrichment values.** ChIP/input enrichment values were calculated over each coding region of the *P. falciparum* 3D7 reference genome from PlasmoDB v26 (www.plasmodb.org) in 3D7/GFP-PfAP2-HC, 3D7/DDGFP-PfAP2-HC ON and OFF Shield-1, 3D7/WT, 3D7/PfAP2-HC-KO, 3D7/GFP-PfAP2-HC-ΔDBD and 3D7/GFP-PfAP2-HC/PfHP1-mScarlet-glmS parasites at 36-44 hpi. Columns A-B: Gene ID and gene annotation. Column C: 3D7/GFP-PfAP2-HC GFP ChIP/input values. Column D: 3D7/GFP-PfAP2-HC HP1 ChIP/input values. Column E: 3D7/DDGFP-PfAP2-HC ON Shield-1 HP1 ChIP/input values. Column F: 3D7/DDGFP-PfAP2-HC OFF Shield-1 HP1 ChIP/input values. Column G: 3D7/WT HP1 ChIP/input values. Column H: 3D7/PfAP2-HC-KO HP1 ChIP/input values. Column I: 3D7/GFP-PfAP2-HC-ΔDBD GFP ChIP/input values. Column J: 3D7/GFP-PfAP2-HC-ΔDBD HP1 ChIP/input values. Column K: 3D7/GFP-PfAP2-HC/PfHP1-mScarlet-glmS GFP ChIP/input values (Experiment 1). Column L: 3D7/GFP-PfAP2-HC/PfHP1-mScarlet-glmS HP1 ChIP/input values (Experiment 1). Column M: 3D7/GFP-PfAP2-HC/PfHP1-mScarlet-glmS GFP ChIP/input values (Experiment 2). Column N: 3D7/GFP-PfAP2-HC/PfHP1-mScarlet-glmS HP1 ChIP/input values (Experiment 2).

**Supplementary Dataset 2. Processed microarray data obtained from 3D7/DDGFP-PfAP2-HC parasites.** Columns A-B: Gene ID and gene annotation. Column C: HP1 target genes[4,5]. Columns D-H: Cy5/Cy3 log2 ratios for all transcripts and five TPs (8-16 hpi, 16-24 hpi, 24-32 hpi, 32-40 hpi, 40-48 hpi) harvested from 3D7/DDGFP-PfAP2-HC parasites cultured in presence of Shield-1 (ON). Columns I-M: Cy5/Cy3 log2 ratios for all transcripts and five TPs harvested from 3D7/DDGFP-PfAP2-HC parasites cultured in absence of Shield-1 (OFF). Columns N-R: fold change in gene expression (log2) in 3D7/DDGFP-PfAP2-HC parasites cultured in absence Shield-1 (OFF) compared to 3D7/DDGFP-PfAP2-HC parasites cultured in presence of Shield-1 (ON) for each of the five paired TPs. Column s: mean fold change in gene expression across all five TPs. Column T: p-value (paired two-tailed Student's t-test). Column U: negative log10 of the p-value.
